# Development of a Fully Non-Viral 1XX-enhanced BCMA CAR-T Cell Therapy for Multiple Myeloma

**DOI:** 10.64898/2026.04.20.719660

**Authors:** Alexis Talbot, Ke Li, Jae Hyun J. Lee, Shanshan Lang, Chang Liu, Nechama Kalter, Zhongmei Li, Yasaman Mortazavi, Niran Almudhfar, Joseph J. Muldoon, Vincent Allain, William Nyberg, Jing-Yi J. Chung, Charlotte Wang, Zhongxia Qi, Netravathi Krishnappa, Alvin S. Ha, Dehui Kong, Derrick Houser, Sreenivasan Paruthiyil, Moloud Ahmadi, Yongchang Ji, Michael Rosenberg, Luis A. Acevedo, Brianna Liang, Kevin Briseno, Serena S. Kwek, Stefanie Bachl, Carter Ching, Julia Carnevale, Petros Giannikopoulos, David Y. Oh, Alexander Marson, Ayal Hendel, Thomas Martin, Justin Eyquem, Brian R. Shy

## Abstract

Multiple myeloma (MM) is a clonal plasma cell malignancy characterized by bone marrow infiltration, monoclonal immunoglobulin production, and microenvironmental dysregulation that leads to systemic organ damage. The advent of B-cell maturation antigen (BCMA)-directed chimeric antigen receptor (CAR) T-cell therapy has induced unprecedented responses and durability for patients with relapsed/refractory MM. These outcomes are rarely observed with prior salvage strategies, although relapse remains the predominant long-term challenge for most patients. The two currently approved BCMA CAR-T cell products use viral vectors to semi-randomly insert the CAR gene, which results in heterogeneous genomic composition and variability in efficacy, safety, and product consistency. To address these challenges, we integrated targeted CRISPR genome engineering with precise CAR transgene insertion at the T-cell receptor alpha constant (*TRAC*) locus, 1XX CAR signaling architecture to enhance potency and durability, and non-viral manufacturing with a single-stranded DNA repair template to improve efficiency and yield. This approach confers physiological CAR expression, reduces insertional mutagenesis, and improves persistence by mitigating tonic signaling and exhaustion. Our GMP manufacturing process consistently achieved high CAR integration (37.7–72.7%) and yields across all full-scale runs and met predefined release criteria for identity, purity, safety, and quality. In NSG mouse models of MM, the UCCT-BCMA-1 product exhibited exceptionally potent tumor control, CAR-T cell expansion 100–1000-fold greater than that of lentiviral constructs, and durable clearance of myeloma cells after multiple rechallenges. These findings establish a CRISPR-edited, fully non-viral manufacturing platform for next-generation 1XX-BCMA CAR-T therapies with enhanced persistence, safety, and efficacy.

**One Sentence Summary:** CRISPR-engineered, *TRAC*-targeted 1XX-BCMA CAR-T therapy with improved safety, potency, and persistence in relapsed and refractory multiple myeloma.

## INTRODUCTION

Multiple myeloma (MM) is a hematologic malignancy characterized by clonal proliferation of malignant plasma cells in the bone marrow, leading to severe complications such as profound anemia, osteolytic bone disease, renal dysfunction, and immunosuppression. Despite therapeutic advances including the development and use of proteasome inhibitors, immunomodulatory agents, and monoclonal antibodies, MM remains largely incurable due to relapse in most patients^1^. For patients with relapsed or refractory (RR) MM, particularly those resistant to multiple lines of therapy, innovative and curative treatment strategies are critically needed.

Chimeric antigen receptor (CAR) T-cell therapy has transformed the treatment landscape of hematologic malignancies, especially with the success of CD19-targeting CAR-T cell therapies against B-cell lymphomas and acute lymphoblastic leukemia^2,3^. For MM, B-cell maturation antigen (BCMA) has emerged as a promising therapeutic target due to its restricted expression on healthy and malignant plasma cells. BCMA-directed CAR-T cell therapies have demonstrated unprecedented response rates in clinical trials for RRMM^4,5^. The Food and Drug Administration (FDA) approvals of idecabtagene vicleucel and ciltacabtagene autoleucel were driven by the KarMMa and CARTITUDE studies, reporting overall response rates of 80–97%^6^.

Despite these remarkable outcomes, several critical challenges persist, including high relapse rates, treatment-related toxicities, and logistical hurdles associated with CAR-T cell manufacturing and accessibility. A substantial proportion of patients experience disease progression after CAR-T cell therapy due to poor CAR-T cell persistence, T cell exhaustion, and in more rare cases, antigen escape, unlike with BCMA-directed bispecific antibodies^7^. In contrast to CD19, loss of BCMA appears to be relatively uncommon, as ∼93% of tumors remain BCMA^+^ at the time of relapse, which suggests T cell dysfunction or loss is a primary failure mode^4^. Additionally, CAR-T-cell–associated toxicities, such as cytokine release syndrome (CRS) and immune effector cell-associated neurotoxicity syndrome (ICANS), and late neurologic and gastrointestinal toxicity, remain significant clinical concerns. The use of lentiviral and retroviral vectors for CAR gene insertion introduces semi-random genomic integration, which increases product heterogeneity and risk of insertional mutagenesis potentially leading to secondary malignancies^9^. Moreover, the requirement for multiple GMP batches of lentivirus introduces variable quality, high cost, and complexity to the supply chain, with disruptions delaying or preventing patient access^8^.

To address these limitations, we combined recent advances in targeted Clustered Regularly Interspaced Short Palindromic Repeats (CRISPR) engineering, CAR design, and non-viral GMP manufacturing to develop a BCMA CAR-T cell therapy with enhanced potency, persistence, and safety. Following benchmarking against current FDA-approved constructs, we identified an optimal CAR architecture incorporating a bb2121 single-chain variable fragment (scFv), CD28 costimulatory domain, and a mutated CD3ζ signaling domain encoding a single immunoreceptor tyrosine-based activation motif (ITAM) termed 1XX^9^. The 1XX domain calibrates T cell activation strength, thereby enhancing potency and durability while minimizing T cell exhaustion and toxicity^9^. A recent first-in-human Phase I study conducted at Memorial Sloan Kettering Cancer Center incorporated the 1XX domain for a gammaretroviral CD19-directed CAR-T–cell product and demonstrated a 75% complete response rate in patients with relapsed or refractory large B-cell lymphoma at a low dose (25×10 CAR-T cells) and with an improved safety profile (∼4% grade ≥3 CRS and ∼7% ICANS) compared to current commercial therapies^10^.

To achieve precise and stable CAR gene integration, we used CRISPR–Cas9–mediated knock-in at the T cell receptor alpha constant (*TRAC*) locus. This strategy both disrupts the endogenous TCR and places the CAR under physiological transcriptional control, which ensures uniform expression and minimizes tonic signaling. Upon antigen engagement, *TRAC*-driven CAR expression couples receptor internalization to regulated re-expression, thereby providing periods of functional rest that limit exhaustion and enhance antitumor efficacy^11,12^. For GMP manufacturing, we adapted our fully non-viral approach that uses hybrid ssDNA templates to improve knock-in efficiency while minimizing cellular toxicity^13^. The final product demonstrates durable clearance of tumor cells at doses ∼10-fold lower than that of current FDA-approved lentiviral products.

Finally, we include our pre-clinical qualification, characterization, safety, and efficacy studies, which supported successful IND clearance for a currently enrolling clinical trial treating patients with RRMM at the University of California, San Francisco (NCT07340853). Manufacturing SOPs and relevant study reports will be made freely available for academic use as a blueprint for investigators wishing to develop similar products at their own institutions.

## RESULTS

### Development of a potent and durable 1XX-enhanced *TRAC* BCMA CAR-T architecture

To identify an ideal BCMA CAR architecture for *TRAC*-targeted integration, we benchmarked several designs in comparison to lentiviral constructs representative of FDA-approved BCMA CAR-T cell therapies: idecabtagene vicleucel (ide-cel; brand name ABECMA), which incorporates the bb2121 scFv binder, 4-1BB costimulatory domain, and WT CD3ζ domain; and ciltacabtagene autoleucel (cilta-cel; brand name CARVYKTI), which includes a tandem VHH1-2 binder, 4-1BB costimulatory domain, and WT CD3ζ domain^4,5^. Knock-in constructs were generated with each binder (bb2121 or VHH1-2), CD28 or 4-1BB costimulatory domains, and WT or 1XX CD3ζ domain (**Figure 1A, Supplementary Figures 1A,2A**)^9^.

**Figure 1:**
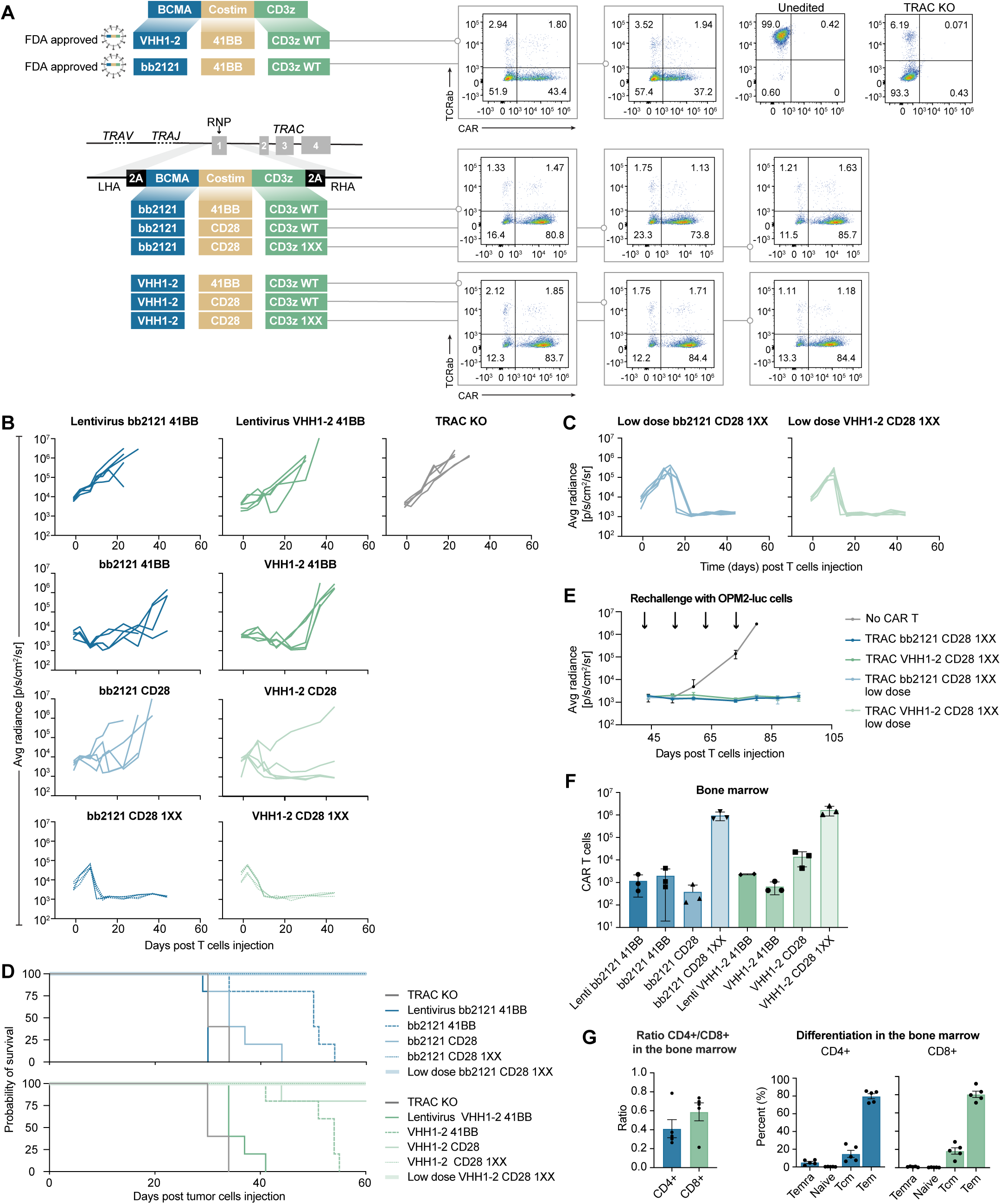
CAR-T engineering: comparison of the efficacy of BCMA CAR-T cells in multiple myeloma model according to binder costimulation domain and CD3ζ. (A) Design of the CAR-T tested: targeted integration of CAR gene at *TRAC* exon 1 via CRISPR/Cas9 using AAV or ssDNA, and random integration using a lentivirus with FDA-approved BCMA CAR (Left). Flow cytometry analysis of CAR and TCR α/ß expression on the surface of non-transduced T cells (T cells), *TRAC* KO cells, and *TRAC* CAR bb2121 or VHH1-2 with 41BB, CD28 WT, or CD28 1XX (Right). (B) Average bioluminescence radiance ((back+front)/2); p/s/cm²/sr) in mice bearing OPM2-luc tumors at the times indicated following IV injection of CAR T-cells (n = 5 mice per group). (C) Average bioluminescence radiance ((back+front)/2); p/s/cm²/sr) in mice bearing OPM2-luc tumors at the times indicated following IV injection of the indicated CAR-T cells with low dose (50,000 CAR-T cells per mouse) (n=5). (D) Overall survival of tumor-bearing mice shown after tumor engraftment following treatment with indicated CAR-T cells (n=5). (E) Average bioluminescence radiance ((back+front)/2); p/s/cm²/sr) in NSG mice treated with CAR-T cells following multiple reinjections of BCMA^+^ OPM2-luc tumors (arrows) (n=5). (F) Analysis of CAR-T cell abundvance in the bone marrow of mice (n=3) at day 10 post T cell injection with the T cell indicated (lentivirus 41BB, *TRAC* 41BB, *TRAC* CD28, and *TRAC* CD28 1XX) bearing either bb2121 or VHH1-2 binder (n=3). (G) Flow cytometry analysis of *TRAC* CAR-T cells in the bone marrow (BM) and spleen, and immunophenotype of CD4^+^ CAR and CD8^+^ CAR present in the bone marrow (n=5).

All constructs underwent efficient knock-in (56.9–88.0%) or lentiviral transduction (37.8–43.0%) in primary human T cells (**Figure 1A**). To facilitate direct comparison with knock-in conditions and prevent xeno-GVHD, CRISPR/Cas9-mediated TCR knockout was performed in addition to lentiviral delivery for the ide-cel and cilta-cel vectors. TCR knock-out was ∼97% across conditions. In vitro cytotoxicity assays demonstrated activity against MM1S and OPM2 myeloma cell lines, with similar performance of all constructs across multiple independent experiments (**Supplementary Figures 1B,2B**).

To benchmark the different CAR designs, we performed an *in vivo* stress test, in which the CAR-T cell dose is lowered to reveal functional limits of different T cell populations. Thus, we used an aggressive OPM2 NSG mouse model and 3×10^5^ CAR^+^ cells per mouse (**Figure 1B,D**). At this dose, both the ide-cel and cilta-cel lentiviral products failed to control tumor proliferation where the knock-in products incorporating the bb2121 or VHH1-2 binders demonstrated initial tumor control, but some mice faced relapse ∼30 days post-infusion. In contrast, the combination of *TRAC* knock-in with the 1XX CD3ζ domain and CD28 co-stimulation demonstrated complete and durable tumor clearance out to >90 days post-infusion for all animals, using either bb2121 or VHH1-2 as the binder (**Figure 1B,D**).

We next tested a further dose reduction to 5×10^4^ CAR^+^ cells per mouse (equivalent to a ∼20×10^6^ cells in humans by allometric scaling) and again observed complete and durable remission across all mice (**Figure 1C**). Finally, we rechallenged each animal with four or five serial injections of increasing doses of OPM2 cells and observed full protection at both the 5×10^4^ and 3×10^5^ CAR-T doses for the *TRAC* 1XX constructs (**Figure 1E, Supplementary Figure 2E)**. The sustained efficacy with 1XX also translated to improved OS, with no tumor-related deaths observed in mice treated with the 1XX *TRAC* CARs regardless of dose or binder (**Figure 1D, Supplementary Figure 2D**).

Quantification of cells in the bone marrow for mice sacrificed at 10 days after T cell injection indicated more than a 1,000-fold increase in CAR^+^ cells for 1XX *TRAC* BCMA CAR constructs (**Figure 1F**). T cells isolated from the bone marrow exhibited a predominantly effector memory or central memory immunophenotype both at 10 days post-infusion and in separate experiments at three months post-infusion following tumor rechallenge **(Figure 1G, Supplementary Figure 2F**). These results are consistent with phenotypic characterization of CD19 1XX CAR-T cells in the context of B-cell lymphoma^10^. CAR-T cells isolated from the spleen for *ex vivo* culture remained capable of proliferating and killing tumor cells in co-culture and cytotoxicity assays (**Supplementary Figure 2G)**. Based on the significantly enhanced potency and durability, the 1XX bb2121 *TRAC* BCMA CAR (UCCT-BCMA-1) was selected for further pre-clinical advancement.

### Developing a GMP-compatible, fully non-viral CAR-T manufacturing process

To generate a GMP manufacturing process for the UCCT-BCMA-1 product, we adapted our previously described fully non-viral CRISPR-Cas9 knock-in approach using hybrid ssDNA homology directed repair (HDR) templates that incorporate Cas9 target sequences (CTS) for improved knock-in efficiency and yield^13^ (**Figure 2A**). T cells were isolated and activated directly from fresh leukapheresis products using CD3/CD28 Dynabeads and DynaCellect magnetic isolation, then electroporated on Day 2 using the Maxcyte GTx instrument with GMP-grade high fidelity SpyFi Cas9, *TRAC*-targeting single-guide RNA (sgRNA), and 1XX BCMA CAR ssDNA HDR template. After electroporation, cells were expanded for five days in G-Rex gas-permeable culture vessels and cryopreserved using a VIAFreeze Quad controlled-rate freezer.

**Figure 2.**
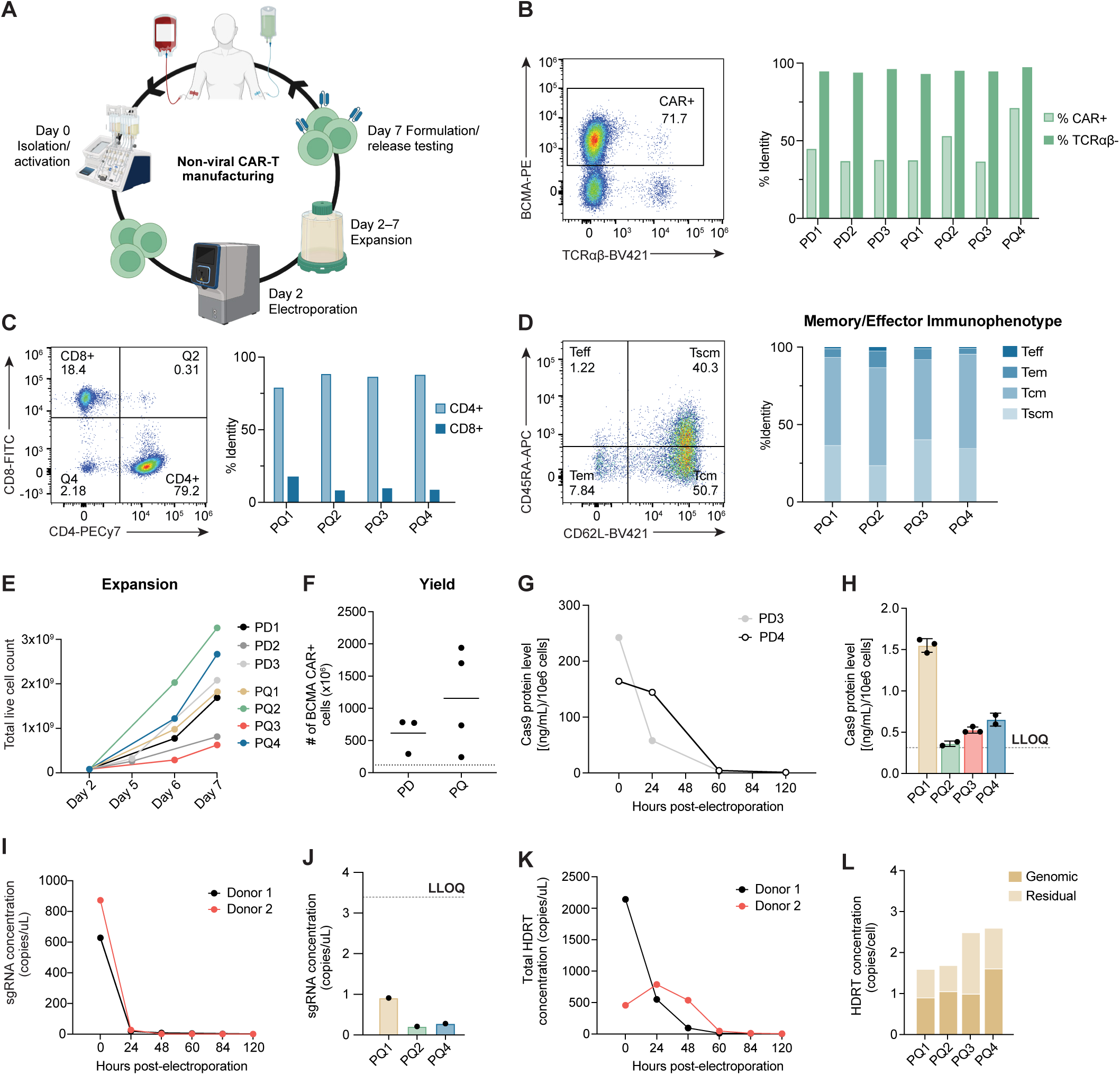
Clinical-scale, non-viral CAR-T cell manufacturing platform. (A) Diagram of optimized non-viral CAR-T cell manufacturing process. On day 0, leukapheresis was processed on DynaCellect using CD3/CD28 Dynabeads to isolate and activate T cells. On day 2, cells were mixed with RNP and ssCTS templates and electroporated using MaxCyte GTx. Cells were then cultured in G-Rex 100M and expanded until day 7. Final drug product was cryopreserved and tested against release criteria before infusion into patients. (B) Representative scatterplot and quantification of BCMA-CAR expression (%CAR^+^) and disruption of TCRα/β expression (%TCRα/β^−^) in fresh drug substance of all full-scale process development (PD) and qualification (PQ) runs. (C) Representative scatterplot and quantification of CD4^+^ and CD8^+^ cells within CD5^+^ live cells in day 7 fresh drug substance of all PQ runs. (D) Representative flow plots for T cell immunophenotype analysis based on CD45RA and CD62L expression. T cell subtypes were defined as naïve/stem cell memory (Tscm, CD45RA^+^/CD62L^+^), central memory (Tcm, CD45RA^−^/CD62L^+^), effector memory (Tem, CD45RA^−^/CD62L^−^), and terminal effector memory (Teff, CD45RA^+^/CD62L^−^). (E) Quantification of T cell expansion during full-scale manufacturing runs. (F) Yield of BCMA CAR-T cells from full-scale manufacturing runs using healthy donors, with dashed line indicating the highest proposed dose level. (G) Residual Cas9 protein in two clinical-scale runs (PD3 and PD4) was shown over time to demonstrate fast clearance of Cas9 post-electroporation. (H) Final drug products of the four process qualification runs had extremely low levels of residual Cas9. Dashed line indicates lower limit of quantification (LLOQ). (I) Concentration of droplets positive for sgRNA detection detected throughout the manufacturing process in two independent donors. A steep reduction in sgRNA levels was observed throughout the manufacturing process, demonstrating effective clearance with an estimated half-life of 4.6 hours. (J) All final drug products showed low levels of residual sgRNA below the established lower limit of quantification (LLOQ) of 3.39 copies/uL indicated by dashed line. (K) Time-course study of total HDRT copy number during manufacturing process, showing 2 to 3-log reduction in template levels from electroporation to harvest and demonstrating effective clearance. (L) Quantification of total and genomic (gray) HDRT copies per cell in PQ drug products with estimated residual HDRT copy number in pink.

Following process optimization, we completed three full-scale Process Development (PD) runs and four Process Qualification (PQ) runs within the GMP environment at the UCSF Human Islet and Cellular Transplantation Facility (HICTF). Flow cytometry analysis demonstrated BCMA CAR knock-in efficiencies ranging from 37.7–72.7% and TCR knock-out >95% (**Figure 2B**). UCCT-BCMA-1 drug products demonstrated an increased CD4^+^:CD8^+^ ratio and predominant stem cell memory and central memory population, an immunophenotype associated with improved self-renewal, persistence, and antitumor activity (**Figure 2C,D**)^14,15^. Each of the seven full-scale manufacturing runs showed cytokine-dependent expansion and T cell purity >99%, yielding 2.4×10^8^–1.9×10^9^ CAR^+^ cells per 1×10^8^ input T cells (**Figure 2E,F, Supplementary Figure 3A,H**). PQ drug products met all proposed release specifications in alignment with FDA recommendations (**Supplementary Table 1**).

To further characterize product safety and support FDA requirements, we performed additional impurity testing for critical genome editing reagents and IL-7/IL-15 cytokines. To assess Cas9 clearance, a quantitative ELISA was developed with a lower limit of quantification (LLOQ) of 0.313 ng/mL. Cas9 levels declined rapidly over time across two full-scale PD runs, reaching undetectable or near-background levels by Day 7 (**Figure 2G**). Final drug product samples from all four PQ runs confirmed minimal Cas9 retention at the time of cryopreservation (**Figure 2H**). Sensitive droplet digital PCR (ddPCR) assays for detection of residual sgRNA and HDR template similarly demonstrated rapid clearance with a half-life of approximately 4.6 hours for sgRNA and 1.1 days for HDR template (**Figure 2I–L**). Residual cytokine levels assessed using a validated Luminex assay demonstrated a 3 to 4 log reduction with final values near or below the assay’s limit of detection and well below physiological ranges (**Supplementary Figure 3B,C**). Altogether, process impurities demonstrated no discernible impact on product safety or activity.

### Off-target analysis and genome integrity

On-target integration of the 1XX BCMA CAR construct at the *TRAC* locus was confirmed using ddPCR and generally correlated with flow cytometry readouts for surface CAR expression (**Figure 2B**, **Figure 3A**). Targeted Locus Amplification (TLA) sequencing further demonstrated precise, site-specific integration of the BCMA CAR transgene at the *TRAC* locus with no evidence of off-target insertions (**Supplementary Report 1**). In line with FDA guidance, potential off-target sites were nominated by multiple orthogonal methods including *in silico* prediction using the COSMID algorithm with up to three mismatches and one DNA or RNA bulge (54 sites), empirical nomination by high sensitivity GUIDE-seq in Cas9 expressing HEK293 cells (157 sites), and GUIDE-seq in three primary human T cell donors using the representative manufacturing process (46 sites) (**Figure 3B, Supplementary Figure 3D,E**)^16–19^. By integrating the in silico and GUIDE-seq lists, we nominated a total of 217 potential off-target sites to evaluate.

**Figure 3:**
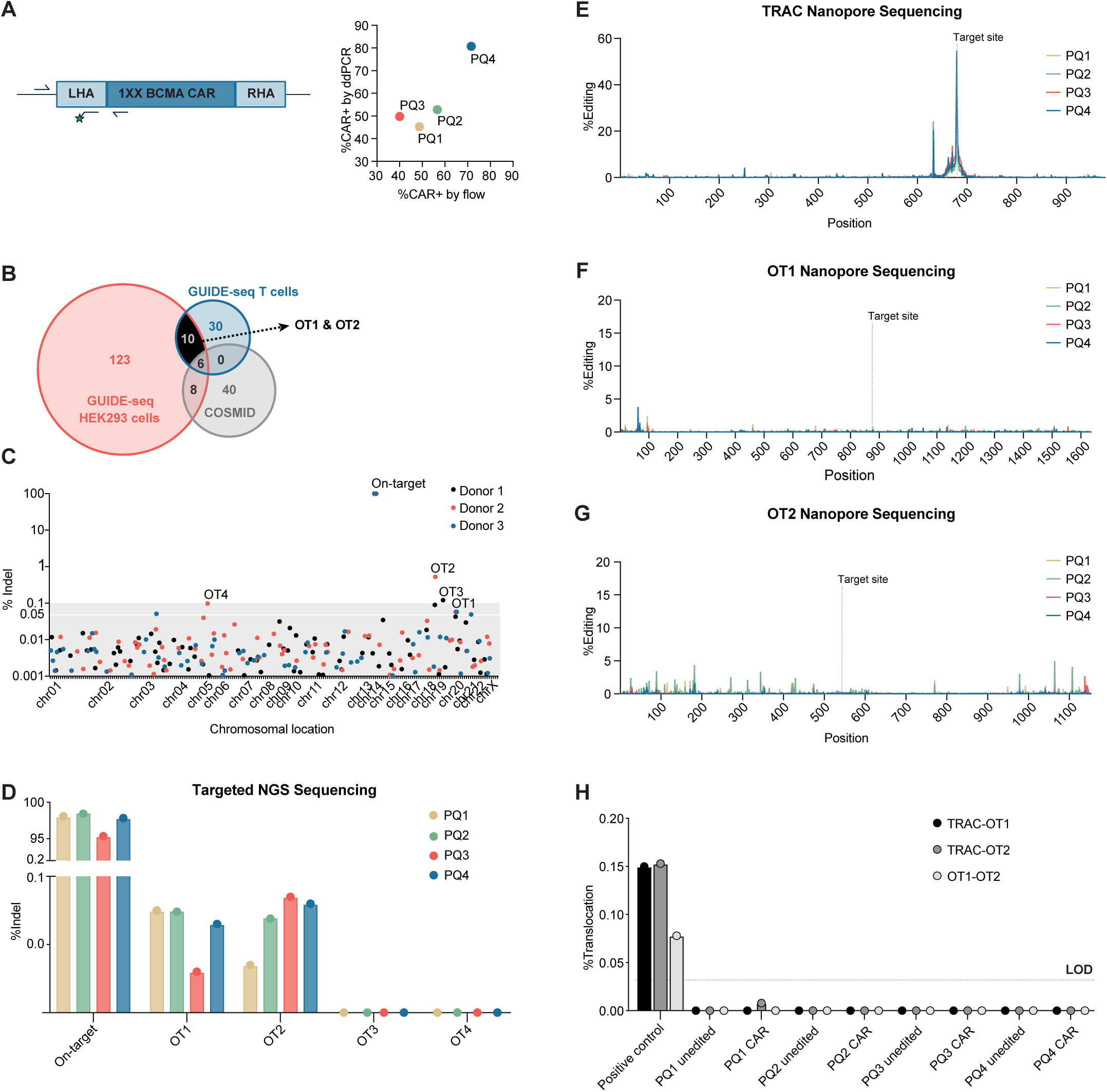
Off-target analysis and genome integrity. (A) Schematic representation of ddPCR assay design for detection of CAR knock-in at the *TRAC* locus. Primer pairs were designed to span the genomic-HDR junction, and FAM-labeled knock-in-specific probe was used to detect genomic integration. CAR knock-in efficiency was independently assessed by flow cytometry based on surface CAR expression and reported as the percentage of CAR-positive cells. ddPCR-based knock-in percentage correlated with flow cytometry-derived CAR expression. (B) Venn diagram of potentially off-target sites nominated by GUIDE-seq in HEK293 cells and primary T cells, and predictions using COSMID. OT1 and OT2 were identified by GUIDE-seq in both HEK293 and primary T cells, but not by COSMID predictions. (C) Identification of off-target indels by rhAmpSeq pooled amplicon sequencing followed by quantification using CRISPECTOR. Each dot represents the editing efficiency (%Indel) at one of the 217 rhAmpSeq loci in different donors edited by *TRAC* RNP. Data point below 0.001% are not shown. (D) Off-target indels across four PQ runs. Bar graph showing the editing efficiency (%Indel) detected at the on-target site and four potential off-target sites (OT1-OT4) in drug products. Editing activity was detected at OT1 and OT2 loci at consistently low levels while no indels were detected for OT3 and OT4. (E-G) Long-read sequencing of the *TRAC* (E), OT1 (F) and OT2 (G) loci. For each nucleotide, the background frequency observed in the unedited control was subtracted from the corresponding drug-product value (negative values set to zero). The resulting background-corrected frequencies are plotted for each locus with cut sites indicated by vertical dotted lines. (H) Three of the six ddPCR assays to detect possible balanced translocations between the *TRAC,* OT1, and OT2 in PQ drug products demonstrate undetectable to below limit-of-detection (LOD) level of translocation events, with no consistent or elevated frequency in gene-edited products compared to unedited controls. Synthetic gBlock gene fragments containing balanced translocation junctions were used as positive controls.

The 217 candidate loci identified by initial screening were analyzed in T cells from three additional healthy donors using a multiplexed rhAmpSeq panel developed by Integrated DNA Technologies (IDT) via the representative manufacturing process. The *TRAC* locus on-target site demonstrated high efficiency editing with insertions and deletions (indels) detected in 98.9–99.8% of reads across all donors (**Figure 3C**). Potential off-target sites that exhibited either low-level indel frequencies across multiple donors (≥0.05%, OT1) or higher frequencies in any single donor (≥0.1%, OT2-4) were assessed by singleplex targeted amplicon sequencing at multiple time points during manufacturing (days 5, 7, and 10) (**Supplementary Figure 3F**). Among these, two sites (OT1 and OT2) showed reproducible but low-level editing (0.1–0.2%) above background across multiple donors. In contrast, no indels were detected for OT3 and OT4 (**Figure 3C, Supplementary Figure 3F**). Further characterization of four UCCT-BCMA-1 process qualification runs confirmed low-level editing for OT1 and OT2 (∼0.1%) (**Figure 3D**). Oxford Nanopore long-read sequencing demonstrated small indels localized to the expected cut sites with no additional large-scale alterations (**Figure 3E–G, Supplementary Figure 3G**). OT1 and OT2 demonstrated no proliferative advantage and map to intronic regions of the *CST7* and *ADAMTSL5* genes, respectively, which are not associated with known pathogenicity, regulatory function, or disease relevance (**Supplementary Figure 3F, Supplementary Table 2**). Both off-target sites were nominated by each of the GUIDE-seq studies and absent from COSMID predictions, which highlights the sensitivity of these assays and high positive predictive value for primary T cell GUIDE-seq methods (**Figure 3B**).

Finally, a comprehensive cytogenetic assessment was conducted to evaluate the genomic stability of UCCT-BCMA-1 cells following CRISPR-Cas9–mediated editing. Three complementary assays were performed across four PQ manufacturing lots including G-banded karyotyping for detection of gross chromosomal abnormalities, single nucleotide polymorphism (SNP) microarray for high-resolution genome-wide copy number variation analysis, and targeted ddPCR for high sensitivity translocation analysis at known double-stranded break (DSB) sites, including *TRAC,* OT1, and OT2. Karyotype analysis indicated a normal diploid karyotype (46,XX or 46,XY) with no numerical or structural abnormalities (**Supplementary Table 3**). No evidence of chromosomal translocations, deletions, duplications, or aneuploidy was observed in any of the tested lots. Similarly, aCGH studies indicated normal genomic profiles with no pathogenic or likely pathogenic copy number alterations (**Supplementary Table 3**). ddPCR assays were designed to evaluate six balanced translocations between the *TRAC*, OT1, and OT2 loci, and no translocation events were observed above the limit of detection (∼0.2%) across all samples (**Figure 3H**).

In summary, the UCCT-BCMA-1 manufacturing process demonstrated highly efficient and specific *TRAC* locus knock-in with minimal off-target activity and no observed large-scale genome alterations.

### Assessment of UCCT-BCMA-1 functional activity and safety

All UCCT-BCMA-1 drug products demonstrated dose-dependent cytotoxicity in *in vitro* co-culture with OPM2 myeloma cell lines with antigen-specific upregulation of CD69 and secretion of IFN-γ, TNF-α, and IL-2 (**Supplementary Figure 4A,B)**. For a subset of donors, we noted reduced post-cryopreservation potency correlated with loss of viability and cell death 24–48 hours after thaw and decreased antitumor activity in the overnight cytotoxicity assay (**Figure 4A, Supplementary Figure 4C,D**). To improve post-cryopreservation recovery, we evaluated alternative DMSO (5–10%) and HSA concentrations (0.25–10%) and observed optimal recovery and potency at 5% DMSO and 10% HSA with similar performance to fresh cells across multiple donors (**Supplementary Figure 4C–F**). This formulation was selected for subsequent manufacturing (**Figure 4A**).

**Figure 4:**
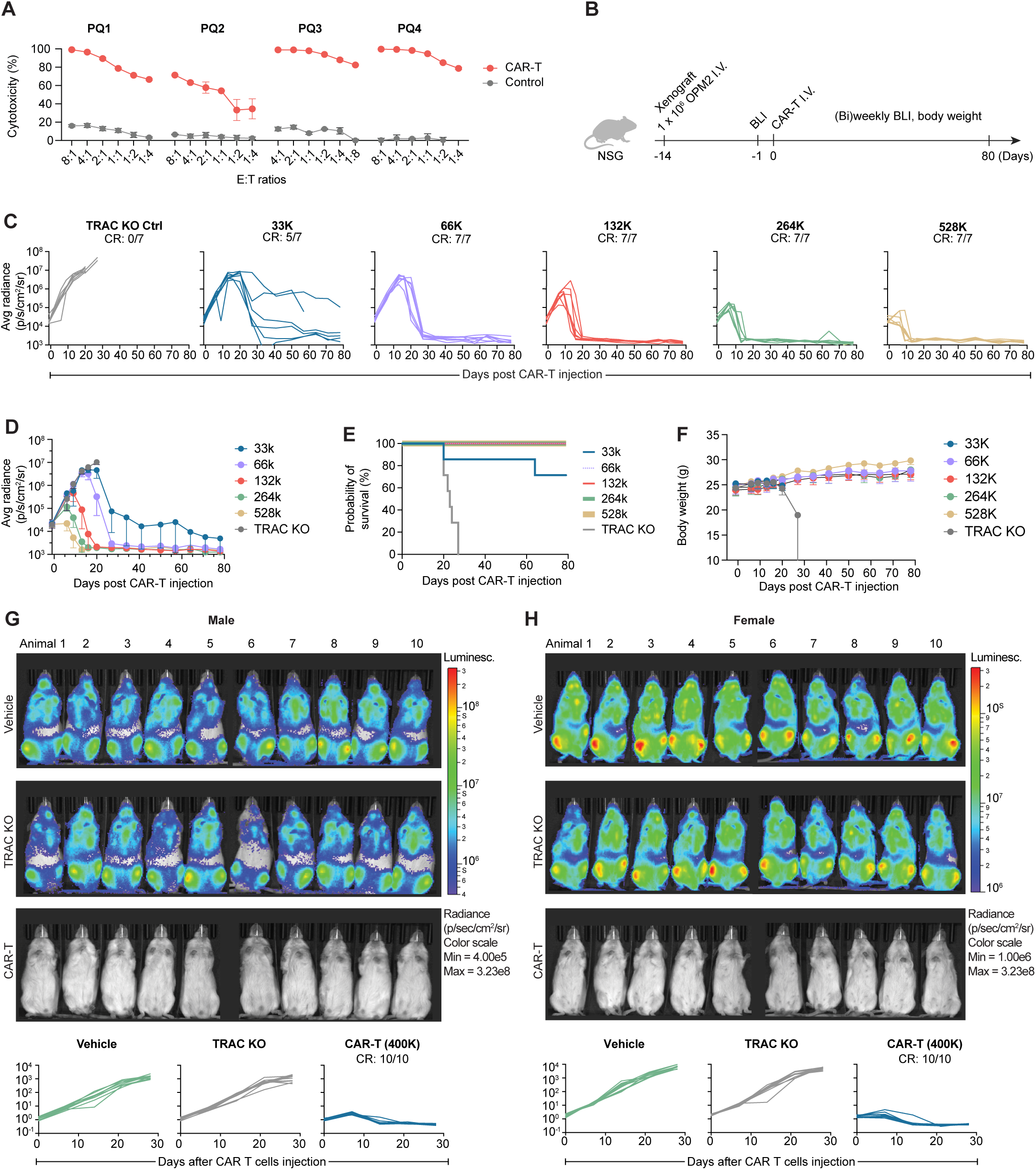
Assessment of UCCT-BCMA-1 functional activity and safety. (A) *In vitro* cytotoxicity was performed by co-culturing the effector (E) UCCT-BCMA-1 cells from PQ runs against the OPM2-luc target (T) cells at the indicated E:T ratios and monitored for target cell killing using the Incucyte live-cell imaging system. PQ3 and PQ4 cells cryopreserved in formulation media containing 10% HSA demonstrate superior post-thaw viability and potency compared to PQ1 and PQ2 cryopreserved in 0.25% HSA. (B) Experimental design to assess the *in vivo* efficacy of PQ drug products in NSG mice inoculated with OPM2-luc myeloma cells two weeks before IV injection of CAR-T cells at indicated doses. BLI indicates bioluminescence imaging. (C) Individual OPM2 growth curves of mice treated various doses of UCCT-BCMA-1 cells from representative run (n=7). CR = complete response. Tumor growth was assessed by BLI in NSG mice bearing OPM2 multiple myeloma xenografts. Radiance units = photons/s/cm^2^/steradian. (D) Average of tumor growth in mice treated with UCCT-BCMA-1 cells product at various doses (n=7). The average luminescence intensity of the tumor for each treatment group was presented as average radiance (p/s/cm^2^/sr). (E) Overall survival of tumor-bearing mice treated with UCCT-BCMA-1 cells at various doses. The Kaplan-Meier curve illustrated the probability of survival for the tumor-bearing mice after receiving treatments at various doses. All treatment groups show significant difference in survival probability compared to the *TRAC* KO control, P = 0.0031 for the 33K dose group and P = 0.0002 for higher dose groups (Log-rank Mantel-Cox test). (F) Overall body weight of tumor-bearing mice treated with UCCT-BCMA-1 cells at various doses. (G-H) An external, independent study assessing tumor growth by bioluminescence imaging (BLI) in NSG mice engrafted with luciferase-expressing OPM2 multiple myeloma cells (n=10). Mice were treated with a single I.V dose of either vehicle, 4×10 *TRAC* KO T cells, or 4×10 PQ3 CAR-T cells. Male (G) and female (H) mice treated with CAR-T cells demonstrated complete tumor control at the end of the study by Day 28, with a significant reduction in luminescent signal compared to both vehicle and *TRAC* KO controls (P < 0.001, Mann–Whitney test). Individual tumor growth curves over the course of the study were quantified in bottom panels.

We next performed *in vivo* dose-finding with two PQ drug products using the OPM2 MM xenograft model at dose levels ranging from 3.3×10^4^ to 5.28×10^5^ per mouse (equivalent to 15×10^6^ to 240×10^6^ cells per adult human based on allometric scaling) (**Figure 4B–F, Supplementary Figure 4H–K**). UCCT-BCMA-1 cells demonstrated potent anti-tumor activity with complete and durable clearance at 6.6×10^4^ cells for PQ3 and 2.6×105 cells for PQ4. Lower doses demonstrated slower kinetics with initial antitumor activity observed ∼10 days post infusion and complete clearance by Day 30, in comparison to clearance by Day 10 for high dose conditions (**Figure 4C,D, Supplementary Figure 4H,I**). CAR-T cell treatment prolonged survival, with all mice (except for two in the lowest dose group) remaining alive and stable for the experiment duration (78 days), whereas controls succumbed by day 20 (**Figure 4E, Supplementary Figure 4J**). Surviving mice exhibited normal weight trajectories and overall good health across all dose levels (**Figure 4F, Supplementary Figure 4K**). Based on these results and prior clinical experience, a dose escalation of 10×10^6^, 30×10^6^, and 100×10^6^ CAR-T cells was selected for the subsequent clinical trial.

A second, independent GLP toxicology study was conducted through a Contract Research Organization (CRO) to assess *in vivo* efficacy, biodistribution, and safety. Mice were randomized by tumor burden into three groups: vehicle control, *TRAC* KO control, and UCCT-BCMA-1 treatment (*n* = 10 per group for male and female mice). Mice received a single I.V. infusion of 4×10^5^ UCCT-BCMA-1 CAR T cells per mouse, which resulted in complete tumor control for both male and female mice by Day 28 (P < 0.001) with no observed toxicity, weight loss, or deaths (**Figure 4G,H, Supplementary Table 4**). Additional safety assessments, including clinical observations, clinical chemistry, hematology, organ weights, and macroscopic pathology, showed no evidence of test article-related toxicity (**Supplementary Table 5**). Histopathology and immunohistochemistry confirmed the presence of human T cells in lymphoid and clearance-associated organs, consistent with T cell trafficking and tumor clearance. No microscopic findings related to UCCT-BCMA-1 were observed.

In summary, the final UCCT-BCMA-1 drug product demonstrated potent and durable anti-tumor activity across all *in vitro* and *in vivo* studies. The product was well tolerated with no evidence of treatment-related toxicity. Animals exhibited stable weight, normal clinical chemistry and hematology, and no macroscopic or histologic signs of organ pathology.

## DISCUSSION

BCMA-targeted CAR-T cell therapy has produced unprecedented response rates in heavily pretreated patients with relapsed and refractory multiple myeloma^4,6^. Nevertheless, despite initial deep responses, most patients ultimately experience disease relapse and progression. In contrast to CD19-directed therapies, relapse following BCMA CAR-T cell treatment is often not associated with antigen loss, as most tumors maintain BCMA expression at progression. These observations point to T cell dysfunction, exhaustion, or loss of persistence as dominant failure modes, rather than target escape. In parallel, treatment-related toxicities, including CRS and neurologic toxicities, as well as manufacturing complexity, cost, and supply-chain constraints, continue to limit the broader impact and accessibility of current commercial products.

We integrated advances in CAR signaling architecture, targeted genome engineering, and fully non-viral GMP manufacturing to address these challenges and develop UCCT-BCMA-1, a BCMA CAR-T cell therapy designed to enhance potency, durability, and safety. Rather than pursuing maximal signaling strength, this work is grounded in the hypothesis that calibrated activation, enforced through both CAR design and genomic context, is a key determinant of long-term therapeutic efficacy. The CD28–1XX–CD3ζ signaling module builds directly on prior mechanistic studies demonstrating that selective attenuation of CD3ζ ITAM signaling can redirect T cell fate toward enhanced persistence and reduced exhaustion^9^. Importantly, this concept has been supported by emerging clinical data in CD19-directed 1XX CAR-T cells, where high response rates were achieved at substantially lower doses with a favorable toxicity profile, providing a strong rationale for extending this architecture to BCMA^10^.

Targeted integration of the CAR cassette into the *TRAC* locus further reinforces this design principle by driving uniform, physiologic CAR expression while eliminating endogenous TCR signaling. This approach reduces tonic signaling, mitigates exhaustion, and decouples CAR activity from variable transgene copy number inherent to semi-random viral integration. Our systematic benchmarking across binders, costimulatory domains, and signaling architectures demonstrates that *TRAC* targeting alone is insufficient to achieve durable tumor control in aggressive myeloma models. Rather, the combination of *TRAC* locus integration with CD28 costimulation and calibrated 1XX signaling emerged as the critical for sustained efficacy.

Functionally, UCCT-BCMA-1 demonstrated exceptional potency and durability *in vivo*, achieving complete and sustained tumor clearance at doses an order of magnitude lower than those typically required for current FDA-approved BCMA CAR-T cell products. The observed *in vivo* expansion, accumulation within the bone marrow, and persistence following serial tumor rechallenge indicate the establishment of functional CAR-T cell memory rather than transient cytotoxic activity. The magnitude of expansion observed, reaching 100–1000-fold higher CAR-T cell numbers as compared with lentiviral constructs, is consistent with a persistence-driven mechanism of action and supports the hypothesis that calibrated signaling enhances long-term fitness rather than acute effector function alone. These properties are particularly relevant in multiple myeloma, where durable disease control likely requires sustained immune surveillance within the bone marrow microenvironment. In addition, effective tumor clearance at low cell doses with gradual *in vivo* expansion may improve the therapeutic index of BCMA CAR-T cell therapies, including the potential to mitigate acute toxicities such as CRS.

Beyond efficacy, genomic safety is a central consideration for engineered T cell therapies, particularly in light of rare reports of CAR-T–associated clonal expansion and T cell lymphoma^20–22^. Despite the rarity of T cell transformation, semi-random viral integration remains an intrinsic limitation of current commercial products, with integration site heterogeneity introducing biological variability and potential long-term genomic risk^23^. At the same time, CRISPR-based genome engineering raises distinct safety considerations, including off-target editing, large-scale genomic alterations, and chromosomal rearrangements arising from double-stranded DNA breaks. In this study, UCCT-BCMA-1 exhibited precise, site-specific integration at the *TRAC* locus with minimal and well-defined off-target editing as assessed by multiple orthogonal nomination and validation strategies. Detected off-target events were rare, low-level, non-coding, and did not demonstrate evidence of clonal expansion or proliferative advantage. Comprehensive genomic analyses, including G-banded karyotyping, genome-wide copy number assessment, targeted translocation assays, and long-read sequencing, revealed no evidence of chromosomal instability, pathogenic rearrangements, or large-scale genomic alterations. Together, these data support the genomic integrity of the UCCT-BCMA-1 manufacturing process.

The fully non-viral manufacturing platform described here also has practical implications for scalability, cost, and manufacturing planning. By eliminating reliance on lentiviral vectors, this approach reduces exposure to batch-to-batch variability, supply constraints, and certain cost drivers associated with viral production. As an illustrative example, the GMP-grade synthetic sgRNA lot purchased for this study (approximately 75 mg total) would be sufficient to support on the order of 500 manufacturing runs at the doses used, while the ssDNA repair template supply (approximately 5 mg total) would support roughly 70 runs. These quantities reflect procurement choices rather than technical limits, and larger reagent lots could be readily obtained if needed, providing predictable and modular starting material for non-viral genome engineering workflows.

Critically, this work extends beyond product development to provide a comprehensive, IND-enabling blueprint for CRISPR-engineered T cell therapies. The manufacturing process, analytical methods, and preclinical safety and efficacy studies were developed in close alignment with current regulatory expectations and supported successful FDA clearance of UCCT-BCMA-1 for a first-in-human Phase I clinical trial at UCSF. Standard operating procedures and supporting study reports will be made available for academic use to facilitate investigator-initiated cell therapy development and enable adaptation of this product design to additional targets and indications.

Several limitations warrant consideration. Although NSG xenograft models provide a stringent and informative assessment of in vivo potency and persistence, they do not fully recapitulate the complexity of the human tumor microenvironment or immune system. Long-term safety, durability, and toxicity ultimately will require validation through clinical follow-up. In addition, although *TRAC*-targeted integration and 1XX signaling appear synergistic in this context, further work will be needed to determine how these design principles generalize across cell types, disease settings, and combinatorial engineering strategies.

In summary, UCCT-BCMA-1 integrates calibrated CAR signaling, precise genome engineering, and non-viral manufacturing to address key limitations of current BCMA CAR-T cell therapies. We demonstrate durable antitumor activity with a favorable safety profile and provide a fully characterized, IND-enabling manufacturing and analytical framework. Our approach can be adapted to other CAR designs and therapeutic targets to support the development of engineered T cell therapies beyond BCMA.

## MATERIALS AND METHODS

### Primary human T Cells

Human peripheral blood mononuclear cells were obtained via leukapheresis from healthy anonymous donors and provided by STEMCELL Technologies (# 70500.1). The mononuclear cells were separated from red blood cells using density gradient centrifugation with LymphoprepTM (StemCell Technologies). Primary human CD3^+^ T lymphocytes cells were isolated and purified using the EasySep Human T cell isolation kit (STEMCELL Technologies #17951) (1:1 beads-to-cell). The isolated T lymphocytes were activated with Dynabeads Human T-Activator CD3/CD28 (ThermoFisher #11141D) in X-Vivo-15 medium (Lonza #BP04-744Q) supplemented with 5% human serum (Gemini Bioproducts #100-512), IL-7 (5 ng/ml: Miltenyi Biotec #130-095-367), IL-15 (5 ng/ml: Miltenyi Biotec #130-095- 760), penicillin-streptomycin (Thermo Fisher, 50 U/ml), and cultured at a density of 1×10^6^ cells per ml. After 2 days of activation, the Dynabeads were removed from the cultured cells using an EasySep Magnet (StemCell Technologies). Medium was exchanged every two to three days, and cells were resuspended at the same density of 1×10^6^ cells per ml.

### Cell Lines

Human MM cell lines positive for BCMA and expressing luciferase, MM1S-luc (t(14;16)(q32;q23)) and OPM2-luc (t(4;14)(p16.3;q32)) (kindly provided by Eric Smith, Dana-Farber), were cultured in RPMI (Gibco) containing 20% fetal calf serum (FCS, Biowest) and 1% penicillin/streptomycin. The cells were split twice a week to maintain a concentration of 1×10^6^ cells/mL. The cells were routinely tested for mycoplasma contamination.

### Guide RNA

We used a previously described guide RNA: (5′-C*A*G*GGUUCUGGAUAUCUGUGUUUUAGAGCUAGAAAUAGCAAGUUAAAAUAA GGCUAGUCCGUUAUCAACUUGAAAAAGUGGCACCGAGUCGGUGCU*U*U*U-3′) that targets the first exon of the constant chain of the TCRα gene (*TRAC*) (Eyquem et al. Nature 2017). This *TRAC* sgRNA was provided by Synthego Corporation (Redwood City, CA) resuspended in the provided Tris-EDTA buffer and stored at -80C°.

### Construction of viral vectors

Lentivirus vectors were produced to express either the single chain variable fragment (scFv) or the tandem nanobody (VHH1-2) CAR with an SFFV promoter. Briefly, the binder is preceded by a truncated CD8α signal peptide and a FLAG-tag and proceeded by a CD8α hinge-transmembrane region, 4-1BB intracellular region, and a CD3ζ signaling region. For lentiviral expression under the 5’ LTR SFFV promoter, the CAR sequences were cloned into pHR_SFFV (addgene #79121) with a P2A sequence and EGFRt downstream of the CAR.

AAV6 vectors were produced to express the scFv or tandem VHH CAR at the *TRAC* locus. Briefly, the binder is preceded by a truncated CD8α signal peptide and a FLAG-tag and proceeded by a CD8α hinge-transmembrane region, 4-1BB intracellular region, and a CD3ζ signaling region. For integrating genes at the *TRAC* locus, homology arms targeting *TRAC* exon 1 were cloned into the AAV plasmid. LHA (610 bp) and RHA (610 bp) sequences are listed in the sequences section below. The genes for the CARs above were cloned in between the homology arms with a P2A sequence and the truncated EGFR (EGFRt) downstream of the CAR.

### AAV production

AAV-ITR plasmids containing homology arms targeting the BCMA CAR and the start of exon 1 of TCRα (*TRAC*) for homology-directed repair (HDR) were used as previously described (Eyquem et al. Nature 2017).The AAV-ITR-containing plasmid was packaged into AAV6 using Polyethylenimine-based co-transfection of HEK293T cells (Polysciences #23966) with pHelper and pAAV Rep-Cap plasmids. Each plate (150 mm) was transfected with 6 µg cargo vector, 8 µg of Rep-Cap plasmid, 11 µg of adenovirus helper plasmid in 200 µl PEI for 72 hours. Transfected 293T cells were collected in AAV lysis buffer (50 mM Tris, 150 mM NaCl) and lysed by three rounds of rapid freeze/thawing, followed by a 1 h incubation at 37 C with 25 units/ml Benzonase (Millipore Sigma #70-664-3). Viral particles were extracted from cells and purified using iodixanol-based density gradient ultracentrifugation (OptiPrep, StemCell Technologies #07820). AV vector titers were determined by qPCR after treating purified samples with DNaseI (NEB #B0303S) and Proteinase K (Qiagen #1114886). qPCR was performed using SsoFast Eva Green Supermix (Bio-Rad#1725201) on a StepOnePlus Real-Time PCR System (Applied Biosystems #4376600) using primers targeting the left homology arm (Forward: CTTTGCTGGGCCTTTTTCCC, Reverse: CCTGCCACTCAAGGAAACCT). Relative quantity was estimated by comparison to a serial dilution of a vector plasmid standard of known concentration.

### Lentiviral production

Lenti-X 293T cells (Takara Bio 632180) were seeded at 18–20×10^6^ cells per 15 cm plate pre-coated with poly-L-lysine 16 hours before transfection and cultured in DMEM + 5% FBS + 1% penicillin-streptomycin. Cells were transfected with lentiviral plasmids and second-generation lentiviral packaging plasmids, pMD2.G (Addgene 12259) and psPAX2 (Addgene 12260), using Lipofectamine 3000 transfection reagent per the manufacturer’s protocol (L3000001). Six hours post-transfection, the transfection medium was replaced with DMEM + 5% FBS + 1% penicillin-streptomycin supplemented with viral booster reagent according to the manufacturer’s instructions (Alstem VB100). Viral supernatants were collected at 24 and 48 hours and centrifuged at 300g for 10 minutes at 4°C to remove cell debris. Lentiviral particles were concentrated using Alstem precipitation solution (Alstem VC100) and stored overnight at 4°C. The virus was centrifuged at 1,500g for 30 minutes at 4°C, resuspended to 100x the original volume in ice-cold PBS, and stored at -80°C until use. For T cell transduction, concentrated lentivirus was directly added to T cells post-electroporation at a 1:25 v/v ratio with with lenti-boost and X-Vivo-15 medium and mixed.

### Viral transduction

T cells were activated using Dynabeads for 48 hours. Then, 2×10^6^ cells were mixed in 20 µL of P3 nucleofection buffer with 3 µL of RNP per well and electroporated (Lonza; program EH-115). T cells were then resuspended in serum-free T cell medium at a density of 2×10^6^ cells/ml accounting for 33% cell death. For AAV transduction, AAV was added at 15 min after electroporation, at specified MOI of 5×10^-4^. After incubating overnight, cells were then passage into serum-containing AAV free T cell medium and expanded as needed.

### Preparation of Ribonucleoprotein (RNP) Complexes

CRISPR RNA (crRNA) and trans-activating CRISPR RNA (tracrRNA) were chemically synthesized (Edit-R, Dharmacon Horizon), resuspended in 10 mM Tris-HCl (pH 7.4) with 150 mM KCl or IDT duplex buffer at a concentration of 160 µM, and stored in aliquots at -80°C. To generate Cas9 ribonucleoproteins (RNPs), the guide RNA was formed by hybridizing crRNA, complementary to the target gene sequence, with tracrRNA, which binds to Cas9. Equimolar amounts of crRNA and tracrRNA were incubated at 95°C for 5 minutes to form the duplex (crRNA). Finally, RNPs were produced by incubating the preformed duplex with HiFi Cas9 Nuclease V3 (Integrated DNA Technologies) at a 3:1 molar ratio (duplex:Cas9) for 10 minutes at room temperature.

### Preparation of HDRT model

Short single-stranded DNA HDRT (Homology-Directed Repair Template) (<200 bp) were synthesized as Ultramer oligonucleotides (IDT). These oligonucleotides were resuspended at a concentration of 100 µM in distilled water and stored at -20 °C until use. Cas9 Target Sites (CTS) sites were incorporated via an additional 5’ sequence added to the base PCR primers. Amplicons were generated using KAPA HiFi polymerase (Kapa Biosystems), purified by Solid Phase Reversible Immobilization (SPRI) bead cleaning, and resuspended in water at a concentration of 0.5–2 µg/µl, measured by light absorbance on a NanoDrop spectrophotometer (ThermoFisher), as previously described. The annealed ssDNA templates were ready for immediate use in electroporation experiments or aliquoted for long-term storage at −20 °C.

### Gene targeting: transfection/nucleofection

After 48 hours of activation, T lymphocytes were detached from CD3/CD28 Dynabeads, and beads were magnetically removed. T cells were transfected by electroporation in P3 buffer (Primary Cell 4D-NucleofectorTM X kit S Lonza #V4SP-3096) with ribonucleoprotein (RNP) using a 4D-Nucleofector 96-well unit (Lonza #AAF-1003S). For each electroporation reaction, the RNP was generated by co-incubating 60 pmol of recombinant Cas9 protein (QB3 MacroLab) with 120 pmol of *TRAC* sgRNA (Synthego CAGGGUUCUGGAUAUCUGU) at 37°C for 15 minutes. The Cas9 ratio was 1:1. Cas9 was mixed with sgRNA in equal volumes, producing an RNP complex. For electroporation, Cas9 was diluted in RNP buffer to 40 µM, and sgRNA was diluted to 60 µM in IDT duplex buffer (30 mM HEPES pH 7.5, 100 mM potassium acetate). For each well, 2×10^6^ cells in 20 µL of P3 buffer were mixed and electroporated with 3 µL of RNP per well using the Lonza EH-115 Nucleofector program. Immediately after electroporation, the cells were diluted by adding 200 µL of serum-free complete medium and incubated at 37°C, 5% CO2 for 30 minutes. The cells were then diluted in medium to 2×10^6^ cells per mL for knock-in experiments. Recombinant AAV6 donor vectors were added to the culture 30 to 60 minutes after nucleofection at an MOI of 5×10^4^ vg/cell, and the culture was incubated overnight. The next day, the AAV-containing medium was replaced with fresh complete medium, and the cells were cultured under standard conditions (37°C, 5% CO2) in complete medium to maintain a density of 1×10^6^ cells per mL every two to three days. Knock-out and knock-in efficiencies were assessed by staining for TCR with an anti-TCRα/β antibody (Miltenyi Biotec) and CAR with Myc-Tag (Cell Signaling Technology) or eGFRt, followed by flow cytometry using a BD LSRFortessa X-50 instrument. Cells were collected 72 hours later for phenotypic characterization and *in vitro* and *in vivo* use.

### Flow cytometry

Cells were stained in 50 µl FACS buffer (2% FBS and 1mM EDTA in PBS) for 20 minutes at 4C at a 1:100 dilution of the following antibodies: AF488 TCRαβ, BV711 EGFRt, CD4, CD8, and FLAG-tag. For CAR detection, T cells were stained with biotin-conjugated recombinant BCMA and secondary PE streptavidin or with EGFRt Tag. Flow cytometry was performed using an LSRFortessa X-50 flow cytometer (BD Biosciences). Flow cytometry data were processed and analyzed using FlowJo software (BD Biosciences).

### *In vitro* cytotoxicity killing assays

For all experiments, effective CAR-T cell count was based on the percentage of CAR-T cell knock in (TCR^−^/EGFRt^+^). To measure CAR-T cell cytotoxicity against OPM2-luc or MM1-S-luc cells, MM1S-luc or OPM2-luc BCMA^+^ myeloma cells (5×10^4^ cells / well) were co-cultured with *TRAC* BCMA CAR-T cells, non-modified T lymphocytes, or *TRAC* KO T cells from blood donors in a 96-well clear-bottom plate (Thermo Fisher Scientific #165306) in X-Vivo-15 culture medium, human serum (5%), penicillin-streptomycin (0.5%), IL-7 (5 ng/ml), and IL-15 (5 ng/ml) with a total well volume of 100 μl. The control for the maximum signal was MM1S-luc or OPM2-luc cells alone, and the minimum signal control was tumor cells with Tween20 (0.1%). T cells were serially diluted to achieve E:T ratios from ranging from 2:1 to 1:64 (triplicate wells at each ratio). After 18 hours of co-culture, 100 μl of d-Luciferin (Goldbio, LUCK-1G) was added to each well (final concentration of 0.375 mg/ml) and luminescence was measured using a GloMax Explorer instrument with software v.3 (Promega). Cytotoxicity for each sample was determined using the formula: 100x(1 − (sample − minimum)/(maximum − minimum)). Values are then normalized using live and dead controls to create percent killing values and plotted in Prism GraphPad.

### Mouse Studies

Mice were used in accordance with ethical guidelines approved by the Institutional Animal Care and Use Committee under protocol AN194345-01E at the University of California, San Francisco (UCSF). Before and during the experiment, mice were maintained under antibiotic prophylaxis continuously (Clavamox 1 g/l). We used 8- to 12-week-old NSG male and female mice obtained from The Jackson Laboratory (#00557) or bred in-house. Mice were housed with a 12-hour light/dark cycle, at a temperature of 20 to 26 °C, and humidity between 30 and 70%. Animals were acclimated for one week before any experiments. Mice were inoculated via tail vein injection with 1×10^6^ MM1S-luc or OPM2-luc cells in 200 μl of RPMI 1640 (Gibco), followed by the indicated doses of *TRAC* BCMA CAR-T cells (resuspended in 200 μl of X-Vivo) by tail vein injection 2 to 3 weeks later. The effective number of CAR-T cells was based on the CAR knock-in as described above. Prior to T cell injection, mice were randomized to achieve similar distributions of tumor load in different experimental groups according bioluminescence analysis carried out the day before. At each imaging session, mice received an intraperitoneal injection of luciferin (15 mg/ml in PBS, 200 μl) 10 minutes before imaging and were anesthetized with isoflurane (Medline). Bioluminescence images were taken twice weekly using the Xenogen IVIS imaging system (Perkin Elmer). Images were analyzed with Living Image software (Perkin Elmer), and regions of interest were drawn around each mouse’s head. Bioluminescence data were expressed as mean radiance (photons/sec/cm^2^/sr). The default imaging exposure was 1 minute, with shorter exposures used for images with saturation at 1 minute. BLI values reported are an average obtained from imaging each mouse in two incidences (ventral and dorsal). Mice were euthanized according to the UCSF protocol if mobility loss, weight loss or other signs of morbidity signs were observed. No limits were exceeded in any experiments. For rechallenge experiments, after CAR-T cell injection, mice received escalating doses of OPM2-luc tumors (2×10^6^ and 5×10^6^ cells) weekly via I.V injection for 4 to 5 rechallenges. A control group of age-and sex-matched mice not receiving CAR-T cells was used to ensure the successful engraftment of these tumor cells. The investigators were not blinded to allocation during experiments and outcome assessment.

### Analysis of CAR-T cells in Mouse Bone Marrow and Spleen

Mice bearing OPM2-luc tumors were euthanized at a predetermined time after BCMA CAR injections (day +10 or at the end of rechallenge experiments). For bone marrow extraction, the long bones of each hind limb were dissected, crushed with PBS in a mortar and pestle, and washed with PBS through a filter. Cells were then centrifuged and treated for 2 minutes with ACK lysis buffer (118-156-721, Quality Biological) to lyse red blood cells then resuspended in 300 µl of FACS buffer. After adding Fc Block (10 μl per sample) (130-092-575, Miltenyi Biotec), cells were stained for CAR (1 μl per sample) and incubated for 30 minutes at room temperature. After washing, cells were resuspended in 2% normal mouse serum (Millipore-Sigma) and Fc block (10 μl) and incubated for 20 minutes at room temperature. Cells were then stained with the appropriate antibody mix (100 µl per tube) and incubated for 45 minutes at room temperature. After staining, cells were washed and resuspended in 300 µl FACS buffer with counting beads (50 μl per sample) (C36950, ThermoFisher Scientific) and analyzed by flow cytometry.

Spleens were mechanically dissociated and then passed through a 70 µm nylon mesh. Cells were then processed as described above for bone marrow cells, but with red blood cells lysed for 10 minutes.

### Statistical Analyses

Statistical analyses were performed using GraphPad Prism or R. The specific test used is indicated in the figure legend, and p-values are shown on the graphs.

### GMP Manufacturing Workflow

On day 0, CD3^+^ T cells were enriched and activated using CD3/CD28 Dynabeads on the DynaCellect automated system (Thermo Fisher) from leukapheresis products from deidentified healthy donors (STEMCELL Technologies), and then cultured at 37°C with 5% CO₂. Unless otherwise specified, cells were cultured in TheraPEAK™ X-VIVO™ 15 medium (Lonza) supplemented with 5% human AB serum (Access Cell Culture) and interleukin-7 (IL-7) and interleukin-15 (IL-15) (Miltenyi Biotec) at 20 and 100 U/mL, respectively. After 48 hours of activation, T lymphocytes were detached from CD3/CD28 Dynabeads, and beads were magnetically removed.

Targeted integration of the BCMA CAR was performed using CRISPR-Cas9-mediated homology-directed repair. 4800 pmol of single-guide RNA was complexed with 1200 pmol of Cas9 protein to form the ribonucleoprotein (RNP) complex. A ssDNA HDR template encoding the CAR was annealed with oligonucleotides to generate short dsDNA Cas9 target sites within the 5’ and 3’ end of the HDRT to form 80 pmol of the template. 80×10 activated T cells were resuspended in MaxCyte electroporation buffer, combined with RNP and HDRT, and electroporated using a G-1000 cuvette on the MaxCyte GTx system (Expanded T Cell 4-2 program). Immediately after electroporation, the cells were rested with warmed cytokine-free medium and incubate at 37°C, 5% CO₂ for 30 minutes. The cells were then transferred to a G-Rex 100M culture vessel and diluted in complete medium to 1×10 cells/mL. The next day, the culture vessel was filled to its 1000 mL capacity for expansion with fresh, complete culture media.

Cells were harvested 7 days after activation, and formulated into freezing bags or cryovials, cryopreserved using controlled-rate freezing, and stored in vapor-phase liquid nitrogen below −150°C until use. For any analysis post-cryopreservation, the cells were thawed in a water bath set to 37°C then diluted at a 1:10 ratio with warmed cytokine-free culture media or thawing media (complete culture media with 1% pulmozyme) and spun down at 200 g for 10 minutes.

### Off-Target Analysis

Potential off-target sites for the *TRAC*-targeting sgRNA were first predicted *in silico* using COSMID^16^, allowing up to three total mutations, including mismatches, insertions, or deletions within the protospacer or NGG PAM sequence, using the hg38 reference genome.

For empirical validation, GUIDE-seq was performed in both HEK293-Cas9 cells and primary human T cells^17^. HEK293-Cas9 cells were electroporated with 10 μM *TRAC* sgRNA and a 34-bp dsDNA tag following the manufacturer’s protocol (Lonza SF solution, 4D-Nucleofector CM-130 program) then harvested 72 hours post-electroporation. Primary human T cells were isolated from leukapheresis products of three independent healthy donors (Stemcell Technologies) using the EasySep human T cell isolation kit and activated with CD3/CD28 Dynabeads at a 1:1 bead-to-cell ratio in complete media. Cells were electroporated with ribonucleoprotein (RNP) complexes composed of wild-type SpCas9 (Aldevron) and *TRAC* sgRNA (Synthego) in the presence of the dsDNA tag using the ExPERT GTx GMP electroporator. Cultures were supplemented with cytokines on days 3, 6, and 10, re-stimulated on day 9, and harvested on day 16. Genomic DNA extraction was performed using the QIAamp DNA Mini Kit (Qiagen). The GUIDE-seq library preparation, sequencing, and analysis were performed as previously described using the guideseq package (v1.0.2)^17^. Untreated controls were used to filter background signals.

Off-target validation and quantification were performed using rhAmpSeq (IDT). A 219-plex panel including COSMID- and GUIDE-seq–predicted sites was designed using the IDT rhAmpSeq designer. Genomic DNA was collected from primary human T cells electroporated with GMP-grade SpyFi Cas9 and GMP-grade *TRAC* sgRNA on days 5, 7, and 10. Library preparation and paired-end 2 × 150 bp Illumina MiSeq sequencing were performed, and editing frequencies were analyzed using CRISPResso2 and CRISPECTOR (v1.0.7) with default parameters^24,25^.

For targeted NGS of the resulting loci from the rhAmpSeq panel, genomic DNA collected on days 5, 7, and 10 was quantified (Qubit 4.0, Thermo Fisher Scientific) and subjected to two rounds of PCR to enrich target regions and incorporate Illumina adapters and indices. Libraries were quality-checked using the Agilent TapeStation and sequenced on a MiSeq v3 kit (2 × 300 cycles) to achieve ≥10,000× coverage per locus. Reads were analyzed using CRISPResso2 to determine the frequency of non-homologous end-joining–induced indels.

Selected on-target and off-target loci were further characterized by long-read nanopore sequencing. PCR amplicons were quantified using a Qubit dsDNA BR assay (Thermo Fisher Scientific), and 200 fmol of DNA per sample was used for library preparation using an Oxford Nanopore Technologies ligation-based library preparation workflow. Libraries were sequenced on a MinION device then processed using the ONT Epi2Me Amplicon Workflow. Reads were aligned to the GRCh38 human reference genome, and insertions, deletions, and substitutions were quantified on a per-position basis.

Genome-wide chromosomal integrity was assessed by cytogenetic analyses. Genomic DNA was isolated using the EZ1 DNA Blood Kit (Qiagen). The microassay analysis was performed using the Infinium Global Diversity Array with Cytogenetics-8 v1.0 BeadChip (Illumina), and the data was analyzed using NxClinical software (BioDiscovery).

For conventional karyotyping, actively dividing cells were cultured in PB-MAX Karyotyping Medium for 72 hours and treated with colcemid (0.5 μg/mL) for 30 minutes at 37°C. Cells were harvested and processed for G-banded metaphase chromosome analysis using standard cytogenetic procedures. Metaphase spreads were analyzed using MetaSystems software, and karyotypes were described according to the International System for Human Cytogenomic Nomenclature.

Balanced chromosomal translocations between the on-target *TRAC* locus and nominated off-target sites were quantified using ddPCR. Synthetic gBlock gene fragments (IDT) containing balanced translocation junctions were used as positive controls. Six ddPCR assays were designed to detect balanced translocations between the on-target site and two off-target loci. Primer and probe sequences are listed in **Supplementary Table 6**. Genomic DNA from UCCT-BCMA-1 qualification runs and donor-matched unedited controls was digested with HindIII. ddPCR was performed using a QX200 ddPCR System (BioRad) following the manufacturer’s protocol, then the data was read on a QX200 Droplet Reader and analyzed with QX Manager software. Translocation frequencies were calculated as the ratio of FAM-positive to HEX-positive droplets (RPP30 reference) and reported as events per genome equivalent.

### Flow Cytometry

Flow cytometry was performed on either the BD LSRFortessa X-50 or Attune NxT flow cytometer (Thermo Fisher), and data were analyzed using FlowJo software. Cells were stained with fluorophore-conjugated antibodies targeting surface markers for identity, purity, immunophenotype, and CAR expression, along with viability dyes, as specified in **Supplementary Table 7**. Unless otherwise indicated, staining was performed in FACS (2% FBS and 1mM EDTA in DPBS) buffer.

### In vitro cytotoxicity study

CAR-T cell cytotoxicity against BCMA⁺ myeloma cells (OPM2-luc or MM1S-luc) was evaluated using a luciferase-based assay. Tumor cells were harvested in log-phase growth, washed, and resuspended in T cell medium. Cells were plated in 96-well plates at 20,000 to 50,000 cells per well depending on the experiment. CAR-T cells were diluted to achieve the desired effector-to-target (E:T) ratios (ranging from 8:1 to 1:64) based on the percentage of CAR⁺ cells, with *TRAC* KO T cells used as donor-matched controls at equivalent total cell numbers. CAR-T or control T cells were added per well in duplicate or triplicate; tumor-only wells received medium as a negative control.

Co-cultures were incubated at 37°C, 5% CO₂ for 18–48 hours. Tumor viability was assessed by adding luciferin substrate (0.375–0.75 mg/mL in DPBS) and measuring luminescence on a GloMax Explorer. Cytotoxicity was calculated as 100 × [1 − (luminescence of analyte well / luminescence of tumor-only well)]. If the cytotoxicity was calculated using 100 x [1 − (sample − minimum) / (maximum − minimum)], the values were normalized to live and dead controls and plotted as percent killing.

### In vivo efficacy study

For all experiments, mice were used in accordance with ethical guidelines approved by the Institutional Animal Care and Use Committee under protocol AN194345-01E at the University of California, San Francisco (UCSF). Female NSG mice (∼9 weeks old; The Jackson Laboratory) were acclimated for one week prior to study initiation and maintained under standard housing conditions. Prophylactic antibiotic water (Clavamox®, 1 g/L) was administered beginning one week prior to tumor inoculation and continued throughout the study. OPM2 multiple myeloma cells (ACC 50, DSMZ) stably expressing eGFP and firefly luciferase under the EF1α promoter were maintained in RPMI 1640 supplemented with 20% heat-inactivated fetal bovine serum and 1% penicillin–streptomycin at 37 °C and 5% CO₂. For xenograft establishment, mice were injected intravenously via tail vein with 1×10 OPM2 cells in 200 µL serum-free, antibiotic-free RPMI 1640. Tumor engraftment was assessed by bioluminescent imaging (BLI) on day 13 post-inoculation. Mice with failed engraftment or extreme tumor burdens were excluded, and remaining animals were randomized into treatment groups. On day 14 post-tumor inoculation, mice received a single intravenous dose (200 µL) of UCCT-BCMA-1 CAR-T cells or control *TRAC* KO T cells resuspended in serum-free RPMI 1640 prior to dosing. CAR-T cell doses were normalized based on CAR^+^ cell frequency and administered across a dose range of 3.3×10 to 5.28×10 CAR^+^ cells per mouse. Donor-matched *TRAC* KO T cells were administered at equivalent total T cell numbers. Tumor burden was monitored by bioluminescent imaging twice weekly for the first two weeks following CAR-T cell infusion and weekly thereafter. Mice were monitored throughout the 80-day study period for body weight, activity, grooming, posture, and signs of distress.

### Residual Process Impurity Assays

Residual cytokines, Cas9, donor template (HDRT), and sgRNA were measured at multiple timepoints during the manufacturing process.

Cytokine measurements were performed on the drug product supernatant by the Human Immune Monitoring Center at Stanford University. The Milliplex HCYTA-60K-PX48 kit (EMD Millipore) was used according to the manufacturer’s instructions, with sample dilutions and the addition of custom Assay Chex control beads (Radix BioSolutions) to all wells. Plates were read on a Luminex FlexMap3D instrument, with duplicate wells for each sample. Data from wells with <20 beads per cytokine were excluded from analysis.

Residual Cas9 was quantified by ELISA. Cell lysate was isolated from cells using cell extraction buffer and sonication. The protein concentration was quantified (Pierce BCA assay, NanoDrop One), then cell lysates were assessed using the Cell Signaling Technologies Cas9 ELISA Kit following manufacturer protocols. Data was read and analyzed on the SpectraMax iD3 instrument.

Residual HDRT was estimated by quantifying total and genomic copies of HDRT using ddPCR. Genomic DNA was purified from cells using the QIAamp DNA Mini Kit (Qiagen), quantified (NanoDrop One), digested with HindIII, and normalized to 10 ng/μL. Primer and probe sequence sets detecting the total HDRT and the genomic CAR integration are listed in **Supplementary Table 6.** ddPCR was performed using a QX200 ddPCR System (BioRad) following the manufacturer’s protocol then the data was read on a QX200 Droplet Reader and analyzed with QX Manager software. The total HDRT and the integrated CAR was calculated as FAM^+^ droplets per cell (normalized to two copies of RPP30) then subtracted to obtain the residual HDRT.

Residual sgRNA was measured by reverse transcription followed by ddPCR. RNA was purified using the Quick-RNA MiniPrep Kit (Zymo) and quantified. 400 ng of the RNA was reverse transcribed using a guide-specific primer then diluted 50,000-fold. Primer and probe sequences are listed in **Supplementary Table 6.** Readout was performed with QX200 Droplet Reader and ddPCR Droplet Reader Oil. Data analysis was conducted with the QX Manager Software, and thresholds were set manually to obtain the number of FAM^+^ droplets.

### Luminex Cytokine Release

Recovered UCCT-BCMA-1 cells and target OPM2 cells were plated in complete medium under the following conditions: donor-matched unedited T cells with and without OPM2 cells, CAR cells with and without OPM2 (effector:target ratio of 1:2), and CAR cells with CD3/CD28 Dynabeads (1:1 bead-to-cell ratio). Cultures were incubated for 48 hours at 37°C, 5% CO₂, then centrifuged at 470 × g for 5 minutes and supernatants collected and stored at −80°C for cytokine analysis. Cytokine concentrations were measured by the Human Immune Monitoring Center at Stanford University as described above.

### Cytokine Independent Growth

Cryopreserved full-scale run products were thawed as previously described then resuspended in pre-warmed complete media for at least 16 hours at 37°C, 5% CO₂ to allow recovery. Following recovery, cells were spun down, resuspended to 1×10 cells/mL and plated in triplicates either with or without IL-7 and IL-15. Cultures were maintained at 37°C, 5% CO₂, and on days 2, 6, and 10, viable cell counts were determined using a NucleoCounter NC-200 with Via2 cassettes. At each time point, cultures that had expanded were resuspended to 1×10 cells/mL with fresh medium to maintain consistent density throughout the assay.

### Charles river Toxicology study

#### Animals and husbandry

Male and female NSG mice (The Jackson Laboratory) were 10 weeks old at study initiation. All animal procedures were conducted by Charles River Discovery Services in accordance with the Guide for the Care and Use of Laboratory Animals, and the facility is AAALAC-accredited.

#### Tumor cell line and xenograft establishment

Firefly luciferase–expressing human OPM2 multiple myeloma cells (DSMZ, ACC-50) were maintained as suspension cultures in RPMI 1640 supplemented with 20% heat-inactivated fetal bovine serum and 1% penicillin–streptomycin at 37 °C with 5% CO₂. Stable luciferase expression was confirmed prior to implantation.

For tumor establishment, OPM2-luc cells were harvested during logarithmic growth, washed, and resuspended in serum-free RPMI 1640 at 5×10 cells/mL. Mice were injected intravenously via tail vein with 1×10 cells in a total volume of 200 µL. Six days after implantation (designated Day 0), tumor burden was assessed by bioluminescent imaging (BLI), and animals were randomized into treatment groups based on flux values.

#### CAR-T cell preparation and dosing

On the day of dosing, cells were thawed, washed, and resuspended in serum-free RPMI 1640. Each mice received a single intravenous infusion (200 µL) of vehicle control, 4×10 *TRAC* KO T cells, or 4×10 CAR^+^ T cells.

#### In vivo bioluminescence imaging

Whole-body BLI was performed on Day 0 and weekly thereafter (Days 7, 14, 21, and 28). Mice received D-luciferin (150 mg/kg, intraperitoneally) and were imaged 10 minutes post-injection under isoflurane anesthesia using an IVIS SpectrumCT system (PerkinElmer). Regions of interest were drawn around each animal, and total photon flux (photons/s) was quantified using Living Image software. Tumor burden inhibition was calculated on Day 28 using median flux values relative to vehicle-treated controls.

#### Sample collection for biodistribution analysis

Blood, bone marrow, and tissue samples were collected on Days 14 and 28. Whole blood was obtained by terminal cardiac puncture under anesthesia and processed for plasma, serum, or whole-blood analysis as appropriate. Bone marrow was collected from bilateral femurs. Major organs (brain, colon, heart, kidneys, liver, lungs, spleen, skin, and gonads) were harvested, weighed, and preserved for downstream molecular or histopathologic analyses. Samples designated for molecular analysis were snap-frozen or preserved in RNA stabilization reagent, while tissues for histopathology were fixed in formalin and processed for paraffin embedding.

#### Quantitative PCR analysis

Genomic DNA was isolated from blood, bone marrow, and tissues using a QIAamp 96 DNA QIAcube HT Kit (Qiagen). DNA was quantified and analyzed by quantitative PCR to determine absolute CAR transgene copy number using a standard curve generated from a linearized plasmid control. Samples were analyzed in triplicate.

#### Cytokine analysis

Plasma samples collected on Day 28 were analyzed for human IFN-γ, TNF-α, IL-2, IL-4, IL-6, IL-10, and IL-17A using a multiplex bead-based immunoassay (MagPlex, Thermo Fisher Scientific) according to the manufacturer’s instructions.

### Tolerability and statistical analysis

Body weight and clinical signs were monitored throughout the study. Animals were euthanized if predefined humane endpoints were reached. Statistical analyses were performed using GraphPad Prism. Differences between treatment groups were assessed using nonparametric tests as indicated, and survival was analyzed using Kaplan–Meier methods.

## Supporting information

Supplementary Report 1

## Acknowledgments

The authors thank Drs. Michel Sadelain and Isabelle Rivière for constructive discussions on the translation of gene edited CAR T cells to the clinic. A. T received financial support from Fondation de France, the Institut Servier, and the ITMO Cancer Program for research training in fundamental and translational cancer research. B.R.S is supported by NIH grants K08CA273529 and L30TR002983, California Institute of Regenerative Medicine (CIRM) grants INFR5-14719 and CLIN1-15060, the UCSF CRISPR Cures for Cancer Initiative, the Stephen and Nancy Grand Multiple Myeloma Translational Initiative, and the UCSF Living Therapeutics Initiative. J.C. received funding from the NIH (K08CA252605), the Parker Institute for Cancer Immunotherapy, the Burroughs Wellcome Fund, California’s Stem Cell and Gene Therapy Agency, CRISPR Cures for Cancer, the V Foundation, the Lydia Preisler Shorenstein Donor Advised Fund, the Pascarella Scholars Fund, and the UCSF Living Therapeutics Initiative.

## Funding

### Author contributions

Conceptualization: AT, KL, JHJL, JE, BRS

Methodology: AT, KL, JHJL, JE, BRS

Investigation: AT, KL, JHJL, SL, CL, NKa, ZL, TM, NA, JJM, VA, WN, JJC, CW, ZQ, NKr, ASH, DK, DH, SP, MA, YJ, MR, LAA, BL, KB, SSK, SB, CC

Visualization: AT, KL, JHJL, JE, BRS

Funding acquisition: JE, TM, AM, BRS

Project administration:

Supervision: JE; BRS

Writing – original draft: AT, KL, JHJL

Writing – review & editing: all authors: AT, KL, JHJL, SL, CL, NKa, ZL, TM, NA, JJM, VA, WN, JJC, CW, ZQ, NKr, ASH, DK, DH, SP, MA, YJ, MR, LAA, BL, KB, SSK, SB, CC, JC, PG, MS, DO, AM, AH, TM, JE, BRS

### Data and materials availability

All data are available in the main text or the supplementary materials.

---

**Supplementary Figure 1**

(A) Schematic CAR integration at the *TRAC* exon 1 locus using CRISPR/Cas9 and design of the various CARs with 2 scFv and 2 costimulatory domains.

(B) *In vitro* cytotoxicity of non-edited T cells, *TRAC* KO cells, and *TRAC* CAR-T cells co-cultured with MM1S-luc cell line at different effector-to-target ratios.

(C) Experimental design to assess the *in vivo* efficacy of different *TRAC* CAR constructs in NSG mice inoculated with OPM2-luc myeloma cells. CAR-T cells were used at the dose of 2e5 cells IV. BLI indicates bioluminescence imaging, OS overall survival

(D) Average bioluminescence radiance ((back+front)/2); p/s/cm²/sr in mice bearing OPM2-luc tumors at the times shown following IV injection of the indicated CAR-T cells (n=5–7).

(E) Survival of mice at the times shown after tumor engraftment following treatment with indicated CAR-T cells (n=5–7).

**Supplementary Figure 2**

(A) Schematic representation of *TRAC* CAR bb2121 constructs with the wild-type CD3ζ or CD3ζ 1XX, CD28 or 4-1BB.

(B) Cytotoxicity assay of the OPM2-luc cell line using non-edited T cells, *TRAC* KO cells or bb2121 CD28 WT, CD28 1XX or 4-1BB *TRAC* CARs at different effector-to-target ratios.

(C) Average bioluminescence radiance ((back+front)/2); p/s/cm²/sr) in mice bearing OPM2-luc tumors at the times shown following IV injection of the indicated CAR-T cells (n=3–7).

(D) Survival of mice at the indicated times after injection with indicated CAR-T cells (n=3–7).

(E) Average bioluminescence radiance ((back+front)/2); p/s/cm²/sr) in mice with bb2121 CD28 1XX CARs (or control group) following reinjections with OPM2-luc cells (arrows) (n=5).

(F) Flow cytometry analysis of *TRAC* CAR-T cells in the bone marrow (BM) and spleen, and immunophenotype of CD4^+^ CAR and CD8^+^ CAR present in the bone marrow (n=5).

(G) Cytotoxicity assay of the OPM2-luc cell line using CAR-T cells isolated from the spleen.

**Supplementary Figure 3**

(A) Manufacturing process yielded highly pure T cells by harvest.

(B-C) IL-7(B) and IL-15(C) concentrations in drug product supernatants were quantified using the validated Luminex assay. Results are presented relative to the cytokine concentrations used during manufacturing (1x and 10-fold serial dilutions thereafter). A 3-4 log reduction in cytokine levels was observed in all process qualification runs, demonstrating effective clearance during manufacturing. Residual cytokine concentrations in the final drug product are predicted to be well below physiological levels following clinical dilution at infusion.

(D-E) Representative sequences of off-target sites identified by GUIDE-seq in primary T cells edited by SpyFi Cas9 and *TRAC* sgRNA (D) and Cas9-expressing HEK293 cells electroporated with *TRAC* sgRNA (E). The intended target sequence is shown in the top line, while the cleaved sites are shown underneath with mismatches to the on-target site shown and highlighted in color. GUIDE-seq sequencing read counts are shown to the right of each site.

(F) Time-course of indel frequency at the on-target and four potential off-target sites in final drug products from one clinical-scale process development run demonstrated low-level indels for OT1 and OT2. No selective advantage was observed for proliferation from Day 5 through Day 10. No indels were detected for OT3 and OT4.

(G) Long-read sequencing near the *TRAC* locus demonstrating localized indel formation centered around the expected cut site. The indel profile was dominated by small insertions and deletions consistent with NHEJ, showing no additional large-scale alterations. The protospacer sequence is underlined and PAM sequence is highlighted in the grey box.

(H) Cytokine-independent growth assay using final drug product of a representative process qualification run showing no cell proliferation in the absence of cytokines.

**Supplementary Figure 4**

(A) The GeoMFI of CD69 expression on CAR^+^ cells or *TRAC* KO T cells after 24-hour co-culturing with or without OPM2 cells showed antigen-specific CD69 expression upon activation.

(B) Normalized supernatant concentrations (pg/ml) of secreted cytokines from UCCT-BCMA-1 T cells, as determined by Luminex assay. CAR-T cells were analyzed in the absence and presence of target OPM2 cell.

(C) Percentage of recovered live cells from cryopreservation media containing 5% DMSO and indicated %HSA immediately post-thaw and after 24- and 48-hours in culture. %Recovery was calculated using (#Recovered viable cells / #Cryopreserved cell x 100%).

(D) Viability of cells cryopreserved in media containing 5% DMSO and indicated %HSA immediately post-thaw and after 24- and 48-hours in culture.

(E) Percentage of recovered live cells from cryopreservation media containing 10% DMSO and indicated %HSA immediately post-thaw and after 24- and 48-hours in culture. %Recovery was calculated using (#Recovered viable cells / #Cryopreserved cell x 100%).

(F) Viability of cells cryopreserved in media containing 10% DMSO and indicated %HSA immediately post-thaw and after 24- and 48-hours in culture.

(G) Comparison of *in vitro* cytotoxicity of CAR-T cells cryopreserved in media containing 0.25% and 10% HSA showing increase potency with increase HSA concentration.

(H) Individual OPM2 growth curves of mice treated various doses of UCCT-BCMA-1 PQ4 product at various doses (n = 7 mice per group). CR = complete response. Tumor growth was assessed by BLI in NSG mice bearing OPM2 multiple myeloma xenografts. Radiance units = photons/s/cm^2^/steradian.

(D) Average of tumor growth in mice treated with UCCT-BCMA-1 PQ4 product at various doses (n=7). The average luminescence intensity of the tumor for each treatment group was presented as average radiance (p/s/cm^2^/sr).

(E) Overall survival of tumor-bearing mice treated with UCCT-BCMA-1 PQ4 product at various doses (n=7). The Kaplan-Meier curve illustrated the probability of survival for the tumor-bearing mice after receiving treatments at various doses. Treatment doses above 256K show significant difference in survival probability in compared to the *TRAC* KO control, P = 0.0001(Log-rank Mantel-Cox test).

