## Supplementary figures and images for "Development of a Fully Non-Viral 1XX-enhanced BCMA CAR-T Cell Therapy for Multiple Myeloma"

### Supplementary Figure 1

Shy and Eyquem, et al, Figure S1

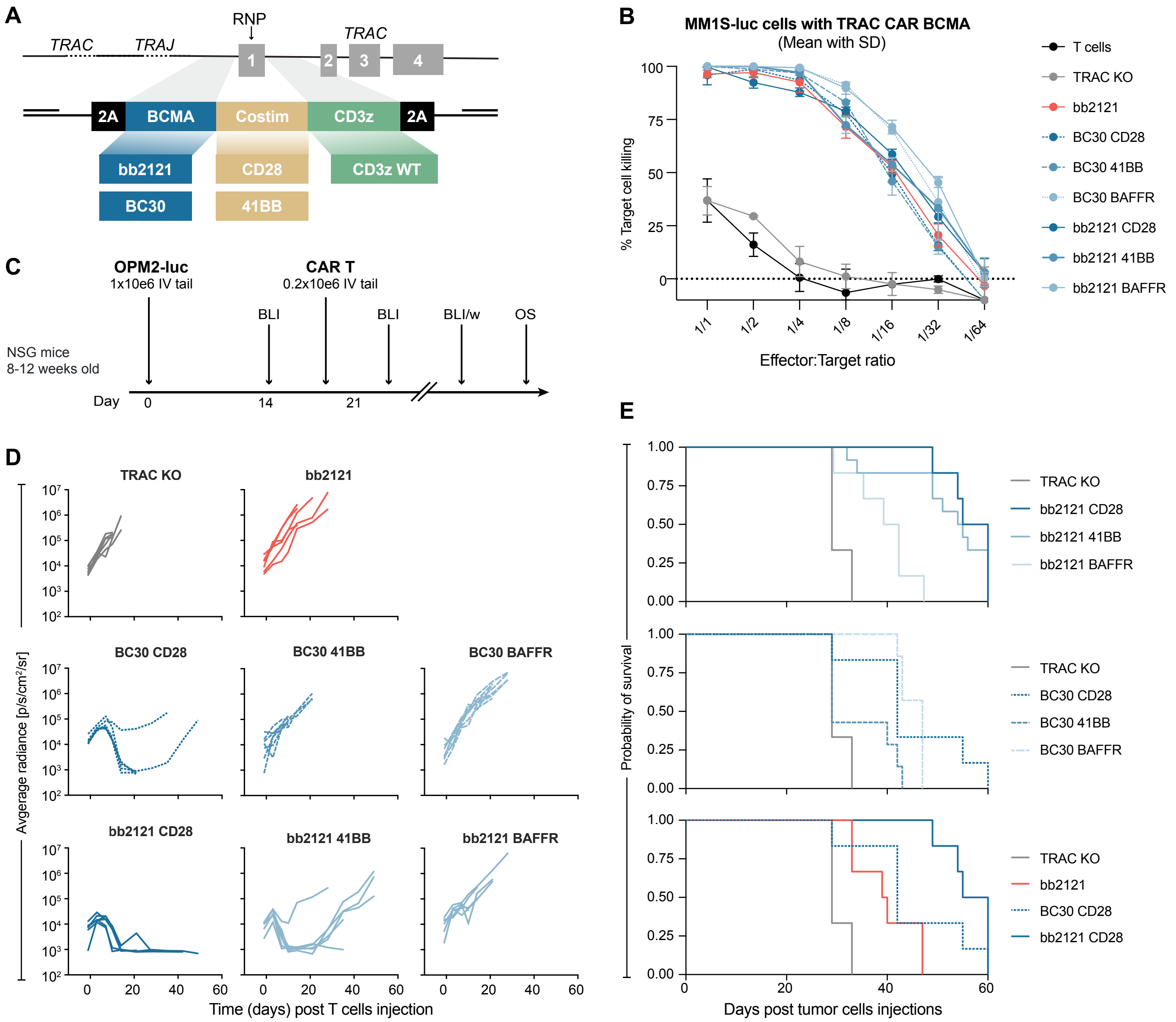


20260225 Shy and Eyquem

### Supplementary Figure 2

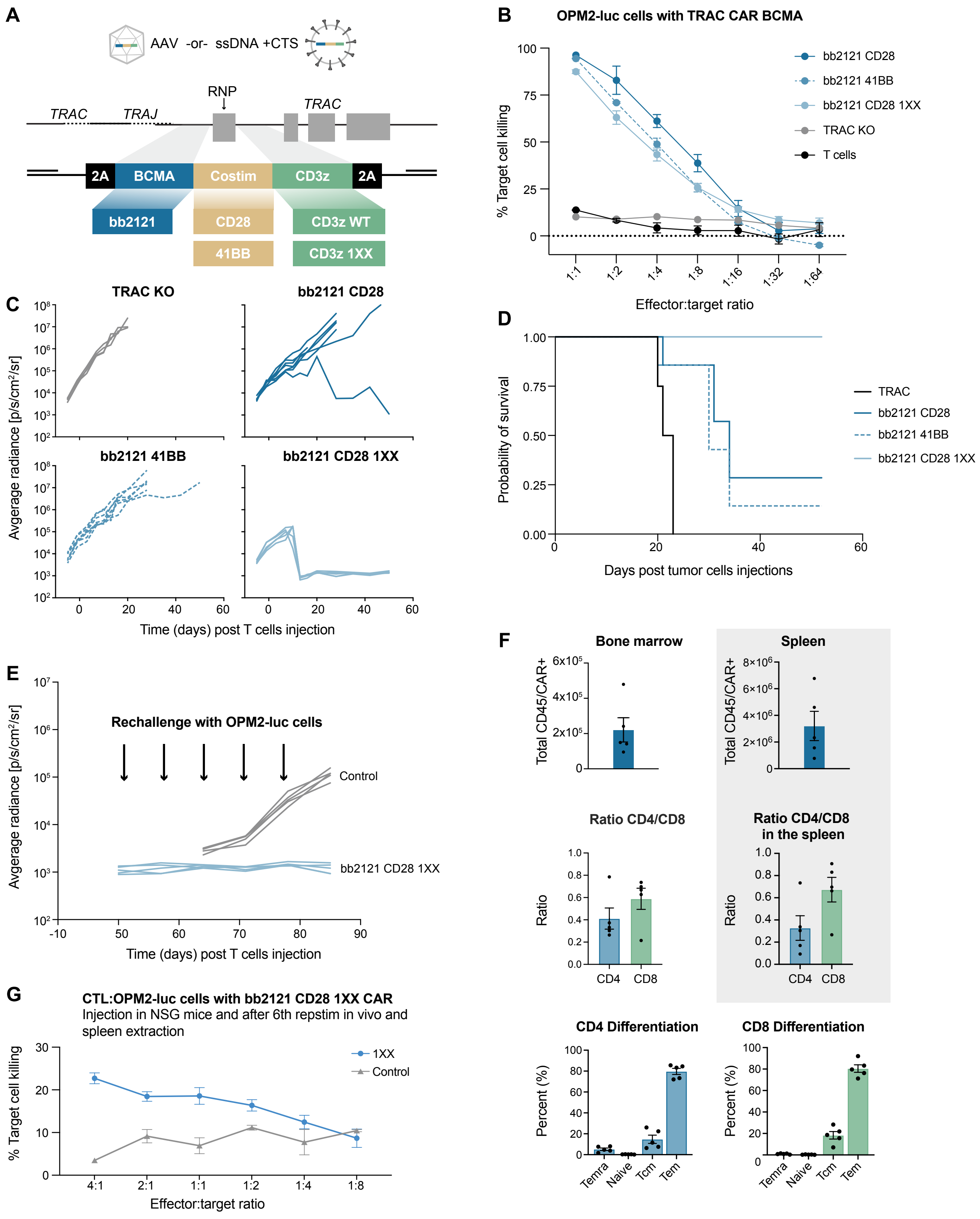

### Supplementary Figure 3

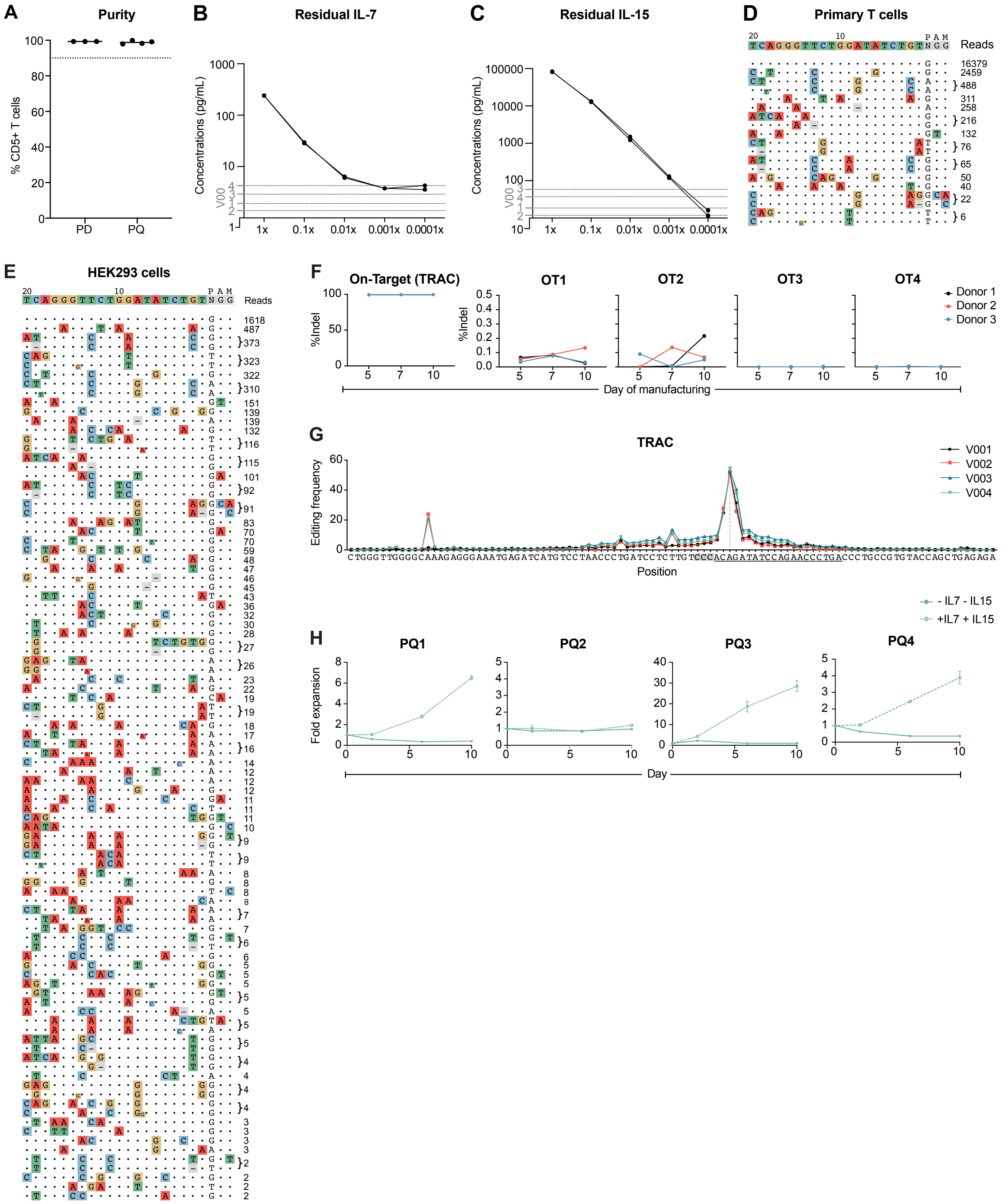

### Supplementary Figure 4

Shy and Eyquem, et al, Figure S4

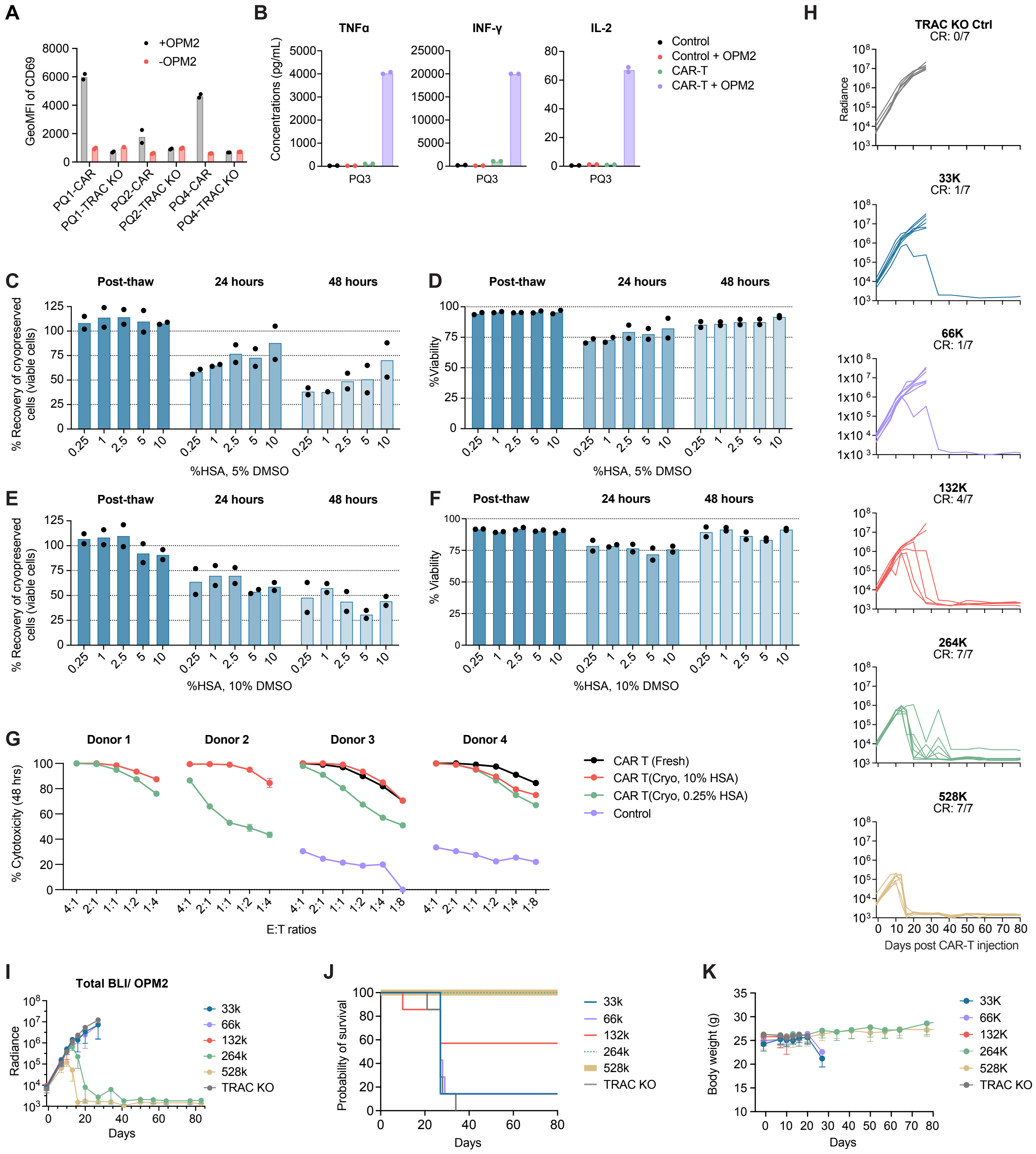
