## Supplementary Report 1 for "Development of a Fully Non-Viral 1XX-enhanced BCMA CAR-T Cell Therapy for Multiple Myeloma"

### Transgene analysis and integration site sequencing of 2 transgenic T-cell lines containing vector hdrt-807-genscript-g526-trac-600

| | |
|---|---|
| Prepared for: | UCSF |
| | 513 Parnassus Avenue, San Francisco, CA94143-0534, US |
| Customer name: | Brian Shy |
| | Clinical Instructor |
| | |
| Internal project number: | 2226 |
| Quote number: | 2021UCSF01/F2022 - 3591 |
| Version: | 2 |
| Date: | 21-Nov-2022 |

CERGENTIS Genetic Engineering

### Goal

In this study, 2 transgenic T-cell lines with the vector hdrt-807-genscript-g526-trac-600 sequence were analyzed.

The aim of this analysis was to:

1. Identify off-target integration events.
2. Estimate the percentage of perfect vs imperfect HR.
3. Report the sequences of the imperfect integration (follow-up study F2022 – 3591)

An overview of the TLA technology and technical details of the performed analyses is provided in the manual "Introduction to the terminology and methods used in TLA analyses_v1".

### Summary

| Sample | Off-Target Events | % perfect HR | % imperfect HR |
|---|---|---|---|
| **Donor1** | 0 | 45 | 7 |
| **Dono2** | 0 | 28 | 1.5 |

### Conclusion

No off-targeted events were observed in either sample. Sample Donor1 was found to have 45% perfect HR and 7% imperfect HR. 28% perfect HR and 1.5% imperfect HR were observed in sample Dono2. The sequences of the imperfect targeting events are provided.

#### TLA, sequencing and data mapping

Viable frozen T-cells were used and processed according to Cergentis' TLA protocol (de Vree et al. Nat Biotechnol. Oct 2014). An overview of the TLA technology and technical details of the performed analyses is provided in the manual "Introduction to the terminology and methods used in TLA analyses_v1".

TLA was performed with 2 independent primer sets specific for the vector sequence (Table 1).

**Table 1**: Primers used in TLA analysis

| Primer set | Name/View point | Direction | Binding position Vector | Binding position genome | Sequence |
|---|---|---|---|---|---|
| **1** | LHA | RV | 302 | chr14: 22,547,166 | CAGCAATATAACTCTGGCAGA |
| | | FW | 383 | chr14: 22,547,227 | TTTCAGGTTTCCTTGAGTGG |
| **2** | CD3z | RV | 1,835 | | CTGGTAATGCTTGCGGGT |
| | | FW | 2,187 | | ACACCTACGACGCCCTTC |
| **3** | TRAC UP | RV | | chr14:22,545,873 | GTGTCAGACTGGAGAAGATC |
| | | FW | | chr14:22,546,111 | CTTCCCAGCAAAGGAACTAT |
| **4** | TRAC DOWN | RV | | chr14:22,549,142 | CTCTTTGCTTTCTCATCCCT |
| | | FW | | chr14:22,549,324 | GGACTCTAGAATGAAGCCAG |

The NGS reads were aligned to the host genome. The human hg38 genome was used as host reference genome sequence.

### Results Donor1 and Dono2

#### Integration sites

##### Whole genome coverage plot

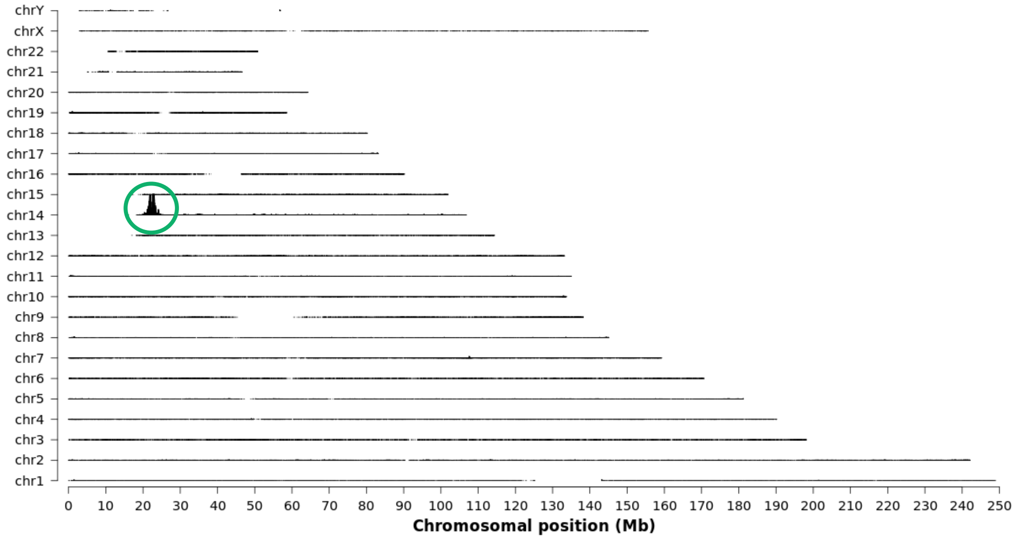


**Figure 1:** TLA sequence coverage across the human genome using vector specific primer set 2 for Donor 1. The chromosomes are indicated on the y-axis, the chromosomal position on the x-axis. Identified integration site is encircled in green.

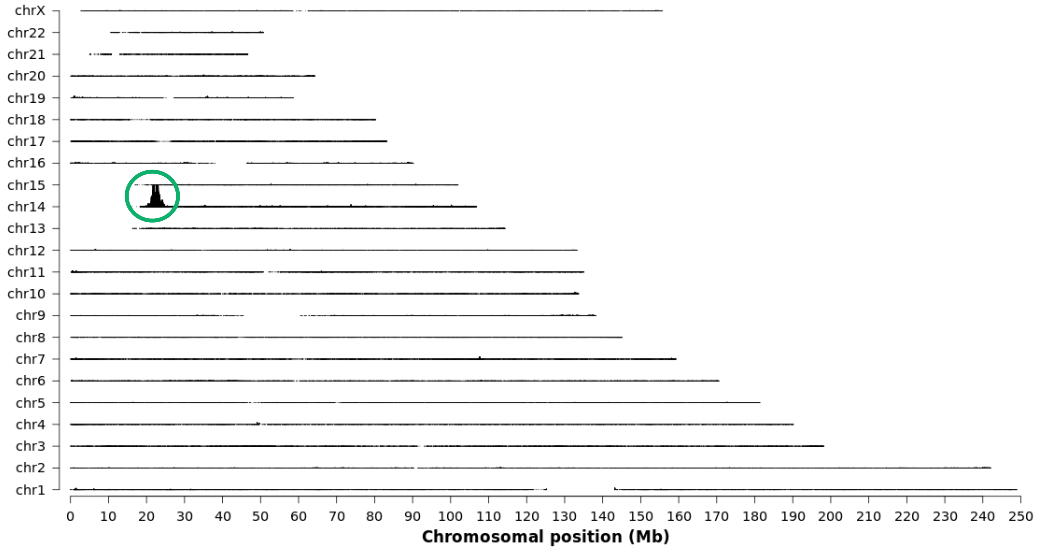


**Figure 2:** TLA sequence coverage across the human genome using vector specific primer set 2 for Dono 2. The chromosomes are indicated on the y-axis, the chromosomal position on the x-axis. Identified integration site is encircled in green.

As shown in figure 1 and 2, the vector has only integrated on chromosome 14 in both samples. No off-target integration events were observed in this sample. Similar results were obtained with primer sets 1, 3 and 4.

#### Targeting events

Based on the counts of the non-targeted reads vs transition of homology arm to insert specific sequence at the LHA the percentage of reads containing an integration event is 52% in sample Donor 1 and 30% in sample Dono2 (table 2 and 3). Only the LHA was evaluated because of a Cas9 cut site in the genome near the inner RHA that causes InDels and skews the data.

Of these integrations for sample Donor1 86% were correctly targeted based on reads containing the ATTCCTGAGATGTAAGGAGCTGCTGTGACT sequence in the vector specific primer set 2 in the vector region 10-61. 14% were incorrectly targeted based on reads containing GATATCTGTCGGAGCTGCTGTGACTTGCTC in the same region (table 4). Combing the results of tables 2 and 4 and thus adjusting the percentage of %LHA for the correct or incorrect targeted outer HA. 45% is perfectly HR and 7% imperfectly HR (table 5) as observed in Figure 3.

Of these integrations for sample Dono2 95% were correctly targeted based on reads containing the ATTCCTGAGATGTAAGGAGCTGCTGTGACT sequence in the vector specific primer set 2 in the vector region 10-61. 5% were incorrectly targeted based on reads containing GATATCTGTCGGAGCTGCTGTGACTTGCTC in the same region (table 4). Combing the results of tables 3 and 4 thus adjusting the percentage of %LHA for the correct or incorrect targeted outer HA. 28% is perfectly HR and 1.5% imperfectly HR (table 5) as observed in Figure 4.

**Table 2**: Percentage of correctly targeted event at inner homology arms for Donor1

| Sample | Read counts LHA | Read counts wildtype | %LHA | % non-targeted |
|---|---|---|---|---|
| **P1** | 860 | 698 | 55 | 45 |
| **P3** | 29 | 15 | 66 | 34 |
| **P4** | 22 | 39 | 36 | 64 |
| **Average** | - | - | 52 | 48 |

Please note the variation with p3 and p4 is most likely due to the location of the primer binding positions on the genome. The numbers were calculating using the LHA. The average is calculated based on averaging the percentages. For non-targeted alleles, the number of reads containing the ATGTCCTAACCCTGATCCTCTTGTC sequence and for LHA the number of reads containing TGTCCTAACCCTGGAATTGGATCCTCTTGTC in region chr14:22,547,464-22,547,545 were counted. Any reads containing SNP or INDEL were excluded.

**Table 3**: Percentage of correctly targeted event at inner homology arms for Dono2

| Sample | Read counts LHA | Read counts wildtype | %LHA | % non-targeted |
|---|---|---|---|---|
| **P1** | 1,333 | 1,556 | 46 | 54 |
| **P3** | 17 | 45 | 27 | 73 |
| **P4** | 36 | 198 | 15 | 85 |
| **Average** | - | - | 30 | 70 |

Please note the variation with p3 and p4 is most likely due to the location of the primer binding positions on the genome. The numbers were calculating using the LHA. The average is calculated based on averaging the percentages. For non-targeted alleles, the number of reads containing the ATGTCCTAACCCTGATCCTCTTGTC sequence and for LHA the number of reads containing TGTCCTAACCCTGGAATTGGATCCTCTTGTC in region chr14:22,547,464-22,547,545 were counted. Any reads containing SNP or INDEL were excluded.

**Table 4**: Percentage of correctly targeted events at the outer left homology arm

| Sample | Read Count correct | Read count incorrect | % Correct | % Incorrect |
|---|---|---|---|---|
| **Donor1** | 720 | 121 | 86 | 14 |
| **Dono2** | 1,585 | 88 | 95 | 5 |

The numbers are calculated based on the breakpoints found aligned to the vector with vector specific primer set 2 at the outer edge of the LHA. For 'correct' targeting the number of reads containing the ATTCCTGAGATGTAAGGAGCTGCTGTGACT sequence and for 'incorrect' the number of reads containing GATATCTGTCGGAGCTGCTGTGACTTGCTC in the vector: 10-61 were counted. Any reads containing SNP or INDEL were excluded.

**Table 5**: Percentage of perfect and imperfect HR

| Sample | % Perfect | % Imperfect |
|---|---|---|
| **Donor1** | 45 | 7 |
| **Dono2** | 28 | 1.5 |

The results are calculated combining the results of table 2 and 3. Adjusting percentage of %LHA for the correct or incorrect targeted outer HA.

**Donor 1**

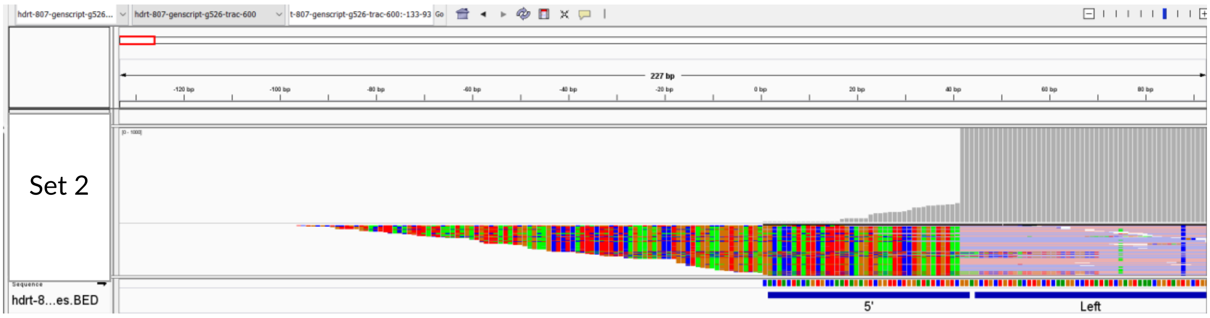


**Figure 3**: TLA sequence coverage (in grey) across part of the left homology arm with vector specific primer set 2. Y-axes are limited to 100x.

The drop of coverage seen in Figure 3 between the homology arms and the 5' CTS indicates perfect HR was observed in this sample. Low coverage across the 5' CTS indicates some imperfect targeting events are present in the studied sample.

#### Dono2

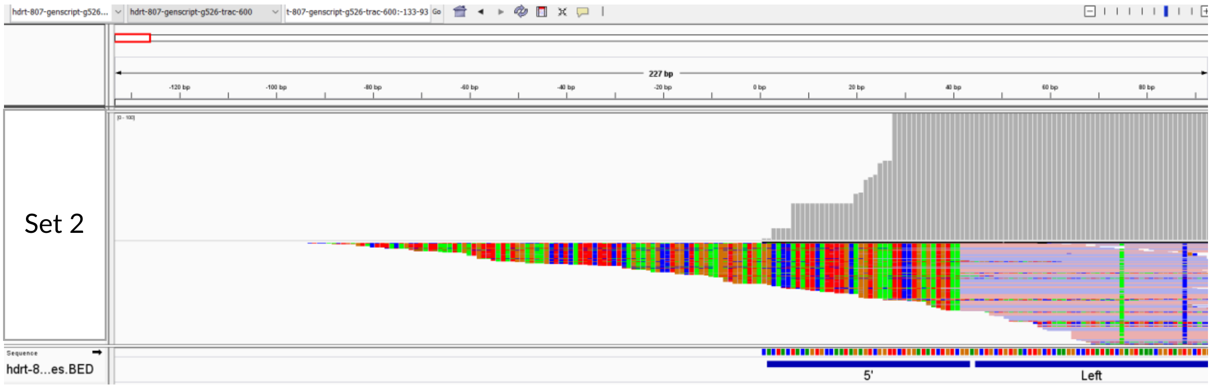


**Figure 4**: TLA sequence coverage (in grey) across part of the left homology arm (top) and right homology arm (bottom) with vector specific primer set 2. Y-axes are limited to 100x.

The drop of coverage seen in Figure 4 between the left homology arms and the 5' CTS indicates perfect HR was observed in this sample. Low coverage across the 5' CTS indicates some imperfect targeting events are present in the studied sample.

#### Sequences of the imperfect targeting events

The sequences and the frequencies of the imperfect targeting events are shown in Table 6 (Donor1) and Table 7 (Dono2).

The breakpoints 1-4 are the same as breakpoints 19-22 but are reported twice since they are mapped at both 5'CTS and 3'CTS. They represent concatemerization of the vector via CTS sequences. Breakpoints 5-8 represent concatemerization of partial vector sequences. Breakpoints 9, 11 and 13 most likely represent the imperfect integration at the targeted locus. Please note that the breakpoints identified with the low number of reads might represent PCR/sequencing artefacts. All described breakpoints are detected with only one primer set therefore independent validation is recommended.

**Table 6**: Breakpoints within CTS sequences in Donor1

| # break point | Breakpoint sequences at 5'CTS | # reads | % of reads |
|---|---|---|---|
| 1 | Vector: 2,933 (tail) fused to Vector: 1 (head)<br>CTTACTGCACTTCTAGGCCTCATTCTAAGCCCCTTCTCCAAGTCCTACAGATATCCAGAACCT<br>ATGCAGGTTCACTACTACAGTGCCAATAGAGTCGGTTCTGGATATCTGTCGGAGCTGCTGTG<br>ACTTGCTCAAGGCCTTATATCGAG | 5 | 3,6 |
| 2 | Vector: 2,916 (tail) fused to Vector: 3 (head)<br>CTCTCCAACTGAGTTCCTGCCTGCCTGCCTTTGCTCAGACTGTTTGCCCCTTACTGCACTTCT<br>AGGCCTCATTCTAAGCCCCCTCTCCAAGTCCTACAGATATCCACTACTACAGTGCCAATAGA<br>GTCGGTTCTGGATATCTGTCGGAGCT | 2 | 1,4 |
| 3 | Vector: 2,910 (tail) fused to Vector: 4 (head)<br>GCCTTTGCTCAGACTGTTTGCCCCTTACTGCACTTCTAGGCCTCATTCTAAGCCCCTTCTCCA<br>AGTCCTACATACTACAGTGCCAATAGAGTCGGTTCTGGATATC | 2 | 1,4 |
| 4 | Vector: 2,910 (tail) fused to Vector: 38 (head) with 6 homologous bases<br>GCCTCAGTCTCTCCAACTGAGTTCCTGCCTGCCTGCCTTTGCTCAGACTGTTTGCCCCTTACT<br>GCACTTCTAGGCCTCATTCTAAGCCCCTTCTCCAAGTCCTACAGATATCTGTCGGAGCTGCT<br>GTGACTTGCTCAAGGCCTTATATCGA | 26 | 18,7 |
| 5 | Vector: 730 (head) fused to Vector: 35 (head) with 4 homologous bases<br>CATGGGGCCGGGATATCTGTCGGAGCTGCTGTGACTTGCTCAAGGCCTTATATC | 13 | 9,4 |
| 6 | Vector: 1,095 (tail) fused to Vector: 28 (head) with 1 homologous base<br>TGATGTAGCAGTTTACTACTGTCTTCAGAGCAGAACCATTCCTTCTGGATATCTGTCGGAGCT<br>GCTGTGACTTGCTCAAGGCCTTATATCGAGTAAACGGTAGTGCTGGGGCTTAGACGCAGGT<br>GTTCTGATTTATAGTTCAAAACCTCTA | 1 | 0,7 |
| 7 | Vector: 1,706 (tail) fused to Vector: 29 (head)<br>CATGTGAAAGGGAAACACCTTTGTCCAAGTCCCCTATTTCCCGGACCTTCTAAGCCCTTTTGG<br>GTGCTGGTGGTGGCTGGATATCTGTCGGAGCTGCTGTGACTTGCTCAAGGCCTTATATCGAG<br>TAAACGGTAGTGCTGGGGCTT | 4 | 2,9 |
| 8 | Vector: 1,626 (tail) fused to Vector: 38 (head) with 4 homologous bases<br>CCGTGTCCAGCGAACAGAAACTTATATCAGAGGAAGACCTTGCAATTGAAGTTATGTATCCT<br>CCTCCTTACCTAGACAATGAGAAAAGCAATGGAACCATTATCTGTCGGAGCTGCTGTGACTT<br>GCTCAAGGCCTTATAT | 8 | 5,8 |
| 9 | Vector: 2,338 (tail) / chr14:22,547,549 (tail) fused to Vector: 36 (head) with 5 inserted bases<br>CCCTGGACCCAATATCCAGAACCCTGACCCTGCCGTGTACCAGCTGAGAGACTCTTGAGCTC<br>TGTCGGAGCTGCTGTGACTTGCTCAAGGCCTTATATCGAGTAAACGGTAGTGCTGGGGCTTA<br>GACGCAGGTGTTCTGATTTATAGTTCA | 1 | 0,7 |
| 10 | chr14:22,620,160 (tail) fused to Vector: 27 (head) with 8 inserted bases<br>TTCCTTCATTTCAACTTTGGTGAATCTGACAATTATGTGTCTTGGAGTTGCTCTTCTCAAGGAG<br>TATCTTTGTGGCATTCTCTGTATTTCCTGAATAGCATCTATTCTGGATATCTGTCGGAGCTGCT<br>GTGACTTGCTCAAGGCCTTATAT | 3 | 2,2 |
| 11 | chr14:22,547,493 (tail) fused to Vector: 35 (head) with 1 inserted base<br>CATGTCCTAACCCTGATCCTGATCTGTCGGAGCTGCTGTGACTTGCTCAAGGCC | 8 | 5,8 |
| 12 | chr14:22,647,005 (head) fused to Vector: 37 (head)<br>CCCAATTCATTAGCAAATCCCACTGGCTCTACCTTTATATATATGTCAAATCCCTGTCGGAGC<br>TGCTGTGACTTGCTCAAGGCCTTATATCGAGTAAACGGTAGTGCTGGGGCTTAGACGCAGGT<br>GTTCTGATTTATAGTTCAAAACCTCT | 2 | 1,4 |
| 13 | chr14:22,546,884 (tail) fused to Vector: 41 (head) | 7 | 5,0 |

| | | | |
|---|---|---|---|
| | TACAGTTTATTAAATAGATGTTTATATGGAGAAGCTCTCATTTCTTTCTCAGAAGAGCCTGGCTAGGAAGGTGGATGAGGCACCATATTCATTTTGCAGGTGAAATTCCTGAGATGTACGGAGCTGCTGTGACTTGCTCAAGGCCTTATAT | | |
| 14 | Unknown* fused to Vector: 18 (head)<br>CATGTAGAGTCGGTTCTGGATATCTGTCGGAGCTGCTGTGAC | 4 | 2,9 |
| 15 | Unknown* fused to Vector: 23 (head)<br>CATGTACAGGAGGACACAGTTATCGGTTCTGGATATCTGTCGGAGCTGCTGTGACTTGCTCAAGGCCTTATATCGA | 35 | 25,2 |
| 16 | Unknown* fused to Vector: 24 (head)<br>CATGCCTCCTTAAACCGGTTCTGGATATCTGTCGGAGCTGCTGTGACTTGCTCAAGGCCTTATATCGAGT | 15 | 10,8 |
| 17 | Unknown* fused to Vector: 25 (head)<br>CATGTACAGGAGGACACAGTTATTGGTTCTGGATATCTGTCGGAGCTGCTGTGACTTGCTCAAGGCCTTATATCGAGTAAACGGTAG | 1 | 0,7 |
| 18 | Unknown* fused to Vector: 38 (head)<br>CATGGATATTTGTCGGAGCTGCTGTGACTTGCTCAAGGCCTTATATCGAGTAAACGGTAGTGCTGGGGCTTAGACGCAGGTGTTCTGATTTATAGTTCAAAACCTCTATC | 2 | 1,4 |
| | Total | 139 | |
| | **Breakpoint sequences at 3'CTS** | | |
| 19 | Vector: 2,933 (tail) fused to Vector: 1 (head)<br>CTTACTGCACTTCTAGGCCTCATTCTAAGCCCCTTCTCCAAGTCCTACAGATATCCAGAACCTATGCAGGTTCACTACTACAGTGCCAATAGAGTCGGTTCTGGATATCTGTCGGAGCTGCTGTGACTTGCTCAAGGCCTTATATCGAG | 3 | 6,7 |
| 20 | Vector: 2,916 (tail) fused to Vector: 3 (head)<br>CTCTCCAACTGAGTTCCTGCCTGCCTGCCTTTGCTCAGACTGTTTGCCCCTTACTGCACTTCTAGGCCTCATTCTAAGCCCCCTCTCCAAGTCCTACAGATATCCACTACTACAGTGCCAATAGAGTCGGTTCTGGATATCTGTCGGAGCT | 2 | 4,4 |
| 21 | Vector: 2,910 (tail) fused to Vector: 4 (head)<br>GCCTTTGCTCAGACTGTTTGCCCCTTACTGCACTTCTAGGCCTCATTCTAAGCCCCTTCTCCAAGTCCTACATACTACAGTGCCAATAGAGTCGGTTCTGGATATC | 1 | 2,2 |
| 22 | Vector: 2,910 (tail) fused to Vector: 38 (head) with 6 homologous bases<br>GCCTCAGTCTCTCCAACTGAGTTCCTGCCTGCCTGCCTTTGCTCAGACTGTTTGCCCCTTACTGCACTTCTAGGCCTCATTCTAAGCCCCTTCTCCAAGTCCTACAGATATCTGTCGGAGCTGCTGTGACTTGCTCAAGGCCTTATATCG | 30 | 66,7 |
| 23 | Vector: 2,909 (tail) fused to Vector: 2,538 (head) / chr14:22,547,749 (head)<br>CAGACTGTTTGCCCCTTACTGCACTTCTAGGCCTCATTCTAAGCCCCTTCTCCAAGTCCTACTTCCAGAAGACACCTTCTTCCCCAGCCCAGGTAAGGGCAGCTTTGGTGCCTTCGCAGGCTGTTTCCTTGCTTCAGGAATGGCCAGGTTC | 8 | 17,8 |
| 24 | Vector: 2,923 (tail) fused to Vector: 24 (tail)<br>TCTCTCCAACTGAGTTCCTGCCTGCCTGCCTTTGCTCAGACTGTTTGCCCCTTACTGCACTTCTAGGCCTCATTCTAAGCCCCTTCTCCAAGTCCTACAGATATCCAGAACCGACTCTATTGGCACTGTAGTAGTG | 1 | 2,2 |
| | Total | 45 | |

* Using the current dataset, it is not possible to elongate the read and identify its nature due to a CATG restrictions site at/close to the breakpoint (CATG sites are used to fragment and ligate DNA during TLA)

The breakpoints 1-3 are the same as breakpoints 21-23 but are reported twice since they are mapped at both 5'CTS and 3'CTS. They represent concatemerization of the vector via CTS sequences. Breakpoints 4-8 and 25-27 represent concatemerization of (partial) vector sequences. Breakpoints 9, 12, 14, 16, 17 and 24 most likely represent the imperfect integration at the targeted locus. Please note that the breakpoints identified with the low number of reads might represent PCR/sequencing artefacts. All described breakpoints are detected with only one primer set, therefore independent validation is recommended.

**Table 7**: Breakpoints within CTS sequences in Dono2

| # break point | Breakpoint sequences at 5'CTS | # reads | % of reads |
|---|---|---|---|
| 1 | Vector: 2,914 (tail) fused to Vector: 8 (head) with 1 homologous base<br>GTTCCTGCCTGCCTGCCTTTGCTCAGACTGTTTGCCCCTTACTGCACTTCTAGGCCTCATTCT<br>AAGCCCCTTCTCCAAGTCCTACAGATATACAGTGCCAATAGAGTCGGTTCTGGATATCTGTC<br>GGAGCTGCTGTGACTTGCTCAAGGCC | 20 | 15,3 |
| 2 | Vector: 2,947 (tail) fused to Vector: 22 (head) with 2 inserted bases<br>ACTTCTAGGCCTCATTCTAAGCCCCTTCTCCAAGTCCTACAGATATCCAGAACCTATGCAGGT<br>TATGTTAGTCAGACGGTGTCGGTTCTGGATATCTGTCGGAGCTGCTGTGACTTGCTCAAGGC<br>CTTATATCG | 9 | 6,9 |
| 3 | Vector: 2,916 (tail) fused to Vector: 30 (head) with 1 homologous base<br>GTTCCTGCCTGCCTGCCTTTGCTCAGACTGTTTGCCCCTTACTGCACTTCTAGGCCTCATTCT<br>AAGCCCCTTCTCCAAGTCCTACAGATATCCTGGATATCTGTCGGAGCTGCTGTGACTTGCTCA<br>AGGCCTTATATCGAGTAAACGGTAG | 1 | 0,8 |
| 4 | Vector: 2,947 (tail) fused to Vector: 1 (head) with 3 inserted bases<br>GAACCTATGCAGGTTATGTTAGTCAGACGGTGCACTACTACAGTGCCAATAGAGTCGGTTCT<br>GGATATCTGTCGGAGCTGCTGTGACTTGCTCAAGGCCTTATATCAAGTAAACGGTAGCGCTG<br>GGGCTTAGACGCAGGTGTTCTGATTTT | 1 | 0,8 |
| 5 | Vector: 2,924 (head) fused to Vector: 41 (head) with 16 homologous bases<br>CGTCTGACTAACATAACCTGCATAGGTTCTGGATATCTGTCGGAGCTGCTGTGACTTGCTCAA<br>GGCCTTATATCGAGTAAACGGTAGTGCTGGGGCTTAGACGCAGGTGTTCTGATTTATAGTTC<br>AAAACCTCTATCAATGAGAGAGCAA | 3 | 2,3 |
| 6 | Vector: 1,146 (tail) fused to Vector: 25 (head) with 4 inserted bases<br>ATGTAGCAGTTTACTACTGTCTTCAGAGCAGAACCATTCCTCGCACATTCGGCGGTGGAACG<br>AAGTTGGAAATCAAGGGCTCAACAAGTGGTTTTGGTTCTGGATATCTGTCGGAGCTGCTGTG<br>ACTTGCTCAAGGCCTTATATCAAGTAA | 1 | 0,8 |
| 7 | Vector: 1,662 (head) fused to Vector: 28 (head)<br>GCCACTGTTACTAGCAAGCTATAGCAAGCCAGGACTCCACCAACCACCACCAGCACCCAAAA<br>GGGCTTAGAAGGTCCGGGAAATAGGGTCTGGATATCTGTCGGAGCTGCTGTGACTTGCTCA<br>AGGCCTTATATCGAGTAAACGGTAGTG | 32 | 24,4 |
| 8 | Vector: 1,644 (head) fused to Vector: 42 (head) with 2 homologous bases<br>ACCACCACCAGCACCCAAAAGGGCTTAGAAGGTCCGGGAAATAGGGGACTTGGACAAAGGT<br>GTTTCGGAGCTGCTGTGACTTGCTCAAGGCCTTATATCGAGTAAACGGTAGTGCTGGGGCTT<br>AGACGCAGGTGTTCTGATTTATAGTTC | 1 | 0,8 |
| 9 | Vector: 253 (head) / chr14:22,547,097 (head) fused to Vector: 3 (head)<br>CATGGGAAAAAGGCCCAGCAAAGCAAACTGTACATCTTCTACTACAGTGCCAATAGAGTCGG<br>TTCTGGATATCTGTCGGAGCTGCTGTGACTTGCTCAAGGCCTTATATCGAGTAAACGGTAGT | 8 | 6,1 |
| 10 | chr14:22,627,688 (tail) fused to Vector: 28 (head) with 6 inserted bases<br>GGGGAATGAGGCCCCTCAGACAGGAATCAGGTCTGCTCACCTTGTTTTGTAGGGATCCTGAA<br>CAAACATACGCCAAAATTGGCAAAGCAGGCGGACGCATCTAGATCTGGATATCTGTCGGAG<br>CTGCTGTGACTTGCTCAAGGCCTTATAT | 4 | 3,1 |
| 11 | chr14:22,565,434 (tail) fused to Vector: 30 (head) with 7 inserted bases<br>CATGTGTCCTAACTCCAGCTCTACCATTAATTAGCCGCAAGACTAACCAAATGGATATCTGTC<br>GGAGCTGCTGTGACTTGCTCAAGGCCTTATAT | 1 | 0,8 |
| 12 | Vector: 2,877 (tail) / chr14:22,548,088 (tail) fused to Vector: 21 (head) with 1 homologous base<br>GGGCAGGAGAGGGCACGTGGCCCAGCCTCAGTCTCTCCAACTGAGTTCCTGCCTGCCTGCC<br>TTTGCTCAGACTGTTTGCCCCTTACTGCACTTCTAGGAGTCGGTTCTGGATATCTGTCGGAGC<br>TGCTGTGACTTGCTCAAGGCCTTATAT | 5 | 3,8 |
| 13 | chr14:22,545,736 (head) fused to Vector: 37 (head)<br>CGACGCCCTTCACATGTGGCTGTTGAAATAAAAATTAGTCAATGTGAAATAAAATGCTTTCAA<br>TTAAAAATGGAAACTTCATTTCAAGAGCTCAACTGTCGGAGCTGCTGTGACTTGCTCAAGGC<br>CTTATATCGAGTAAACGGTAGTGCTG | 6 | 4,6 |

| | | | |
|---|---|---|---|
| 14 | Vector: 2,865 (tail) / chr14: 22,548,076 (tail) fused to Vector: 40 (head) with 3 homologous bases<br>AGAGAAGGTGGCAGGAGAGGGCACGTGGCCCAGCCTCAGTCTCTCCAACTGAGTTCCTGCCTGCCTGCCTTTGCTCAGACTGTTTGCCCCTTACTGTCGGAGCTGCTGTGACTTGCTCAAGGCCTTATATCGAGTAAACGGTAGTGCTGGG | 7 | 5,3 |
| 15 | chr14:22,624,373 (tail) fused to Vector: 22 (head) with 6 inserted bases<br>ATTGTACAAAAATAAGACAAAATTAAATATTTTGTCGGTTCTGGATATCTGTCGGAGCTGCTGTGACTTGCTCAAGGCCTTATATCGAGTAAACGGTAGTGCTGGG | 1 | 0,8 |
| 16 | chr14:22,546,654 (head) fused to Vector: 45 (head) with 3 homologous bases<br>ATGTTAGTTGGAGCCACTGACCCTGCCAGAATATGGCCGTGATAGAGTGCTAGTGAGTCATGCAGAGCCACAGCGTCTATCCTTCACCCTCAGAAAGCAGGAGCTGCTGTGACTTGCTCAAGGCCTTATATC | 2 | 1,5 |
| 17 | chr14:22,546,738 fused to Vector: 44 (head) with 2 homologous bases<br>CATGACTCACTAGCACTCTATCACGGCCATATTCTGGCAGGGTCAGTGGAGCTGCTGTGACTTGCTCAAGGCCTTATATCGAGTAAACGGTAGTGCTGGGGCTTAGACGCAGGTGTTC | 15 | 11,5 |
| 18 | Unknown* fused to Vector: 31 (head)<br>CATGGGATATCTGTCGGAGCTGCTGTGACTTGCTCAAGGCCTTATAT | 2 | 1,5 |
| 19 | Unknown* fused to Vector: 25 (head)<br>CATGGTTCTGGATATCTGTCGGAGCTGCTGTGACTTGCTCAAGGCCTTATATCGAGTAAACGGTAG | 4 | 3,1 |
| 20 | Unknown* fused to Vector: 38 (head)<br>CATGTGTCGGAGCTGCTGTGACTTGCTCAAGGCCTTATATCGAGTAAACGGTAGTGCTG | 8 | 6,1 |
| | Total | 131 | |
| | **Breakpoint sequences at 3'CTS** | | |
| 21 | Vector: 2,914 (tail) fused to Vector: 8 (head) with 1 homologous base<br>GTTCCTGCCTGCCTGCCTTTGCTCAGACTGTTTGCCCCTTACTGCACTTCTAGGCCTCATTCTAAGCCCCTTCTCCAAGTCCTACAGATATACAGTGCCAATAGAGTCGGTTCTGGATATCTGTCGGAGCTGCTGTGACTTGCTCAAGGCC | 20 | 40,0 |
| 22 | Vector: 2,947 (tail) fused to Vector: 22 (head) with 2 inserted bases<br>ACTTCTAGGCCTCATTCTAAGCCCCTTCTCCAAGTCCTACAGATATCCAGAACCTATGCAGGTTATGTTAGTCAGACGGTGTCGGTTCTGGATATCTGTCGGAGCTGCTGTGACTTGCTCAAGGCCTTATATCG | 8 | 16,0 |
| 23 | Vector: 2,916 (tail) fused to Vector: 30 (head) with 1 homologous base<br>GTTCCTGCCTGCCTGCCTTTGCTCAGACTGTTTGCCCCTTACTGCACTTCTAGGCCTCATTCTAAGCCCCTTCTCCAAGTCCTACAGATATCCTGGATATCTGTCGGAGCTGCTGTGACTTGCTCAAGGCCTTATATCGAGTAAACGGTAG | 1 | 2,0 |
| 24 | Vector: 253 (head) / chr14:22,547,097 (head) fused to Vector: 3 (head)<br>CATGGGAAAAAGGCCCAGCAAAGCAAACTGTACATCTTCTACTACAGTGCCAATAGAGTCGGTTCTGGATATCTGTCGGAGCTGCTGTGACTTGCTCAAGGCCTTATATCGAGTAAACGGTAGT | 8 | 16,0 |
| 25 | Vector: 2,943 (tail) fused to Vector: 872 (head) with 2 homologous bases<br>CTAAGCCCCTTCTCCAAGTCCTACAGATATCCAGAACCTATGCAGGTTATGTTAGTCAGAGCTTCTGAAAGTGTCACAATCCTTGGCTCCCACCTGATCCATTGGTACCAACAAAAACCTGGGCAGCCCCCGACGCTTCTCATTCAGTTGG | 5 | 10,0 |
| 26 | Vector: 2,920 (tail) fused to Vector: 667 (head) with 2 homologous bases<br>GTTCCTGCCTGCCTGCCTTTGCTCAGACTGTTTGCCCCTTACTGCACTTCTAGGCCTCATTCTAAGCCCCTTCTCCAAGTCCTACAGATATCCAGAACAGATGGATCTGGAGCAACAAACTTCTCACTACTCAAACAAGCAGGTGACGTGG | 3 | 6,0 |
| 27 | Vector: 2,927 (tail) fused to Vector: 1,467 (head)<br>CTGCTCTTCTAGGCCTCATTCTAAGCCCCTTCTCCAAGTCCTACAGATATCCAGAACCTATGACTTCTGCGCTCTCGATTACTCATACGCTATGGATTACTGGGGCCAGGGCACGTCCGTGACCGTGTCCAGCGAACAGAAACTTATATCA | 4 | 8,0 |
| 28 | Vector: 2,930 (tail) fused to unknown*<br>TCTAAGCCCCTTCTCCAAGTCCTACAGATATCCAGAACCTATGCAGTGGGCCTTTTTCCCATG | 1 | 2,0 |
| | Total | 50 | |

* Using the current dataset, it is not possible to elongate the read and identify its nature due to a CATG restrictions site at/close to the breakpoint (CATG sites are used to fragment and ligate DNA during TLA)

### QC information

#### Sample and Study details

| | |
|---|---|
| Sample receipt date | 04-Nov-2021 |
| Condition of sample at receipt | frozen |
| Start date in the lab | 08-Nov-2021 |
| Sequencing run | Run21-064 |
| Deviations from the protocol | none |
| TLApp version: | 0.5.4.0 |

#### Study Personnel

| | |
|---|---|
| Lab technician | Melinda Aprelia, BSc |
| Data Analyst | Cheryl Dambrot, PhD and Irina Sergeeva, PhD (follow-up) |
| QC Analysis and Report | Irina Sergeeva, PhD and Elaine Wong, PhD |

#### Quality control

The results are independently verified and reviewed and are an accurate and complete representation of the study. TLA processing of cells, NGS sequencing, and data analysis (except for estimation of the targeting efficiency) are ISO/IEC 17025:2017 accredited by the Dutch Accreditation Council RvA, Registration number L671.

| | |
|---|---|
| Scientific approval | Irina Sergeeva, PhD - Scientific Account Manager |
| Date | 21-Nov-2022 |
| Signature | |

#### Version tracking

| Version | Date | Introduced changes |
|---|---|---|
| Version 1 | 02-DEC-2021 | - |
| Version 2 | 21-NOV-2022 | Correction of the sentence “Only the LHA was evaluated because of a Cas9 cut site in the genome near the inner RHA that causes InDels and skews the data” on p.5. Correction of the read number in Table 4 on p.6. Section “Sequences of the imperfect targeting events” is added on p.8 |
